# *Wolffia globosa*–derived nanovesicles as a natural source of bioactive metabolites with neuroprotective potential against methylglyoxal-induced oxidative stress

**DOI:** 10.64898/2026.10.31.762905

**Authors:** Ekkaphot Khongkla, Panitch Boonsnongcheep, Monruedee Srisaisup, Theptharin Charuraksa

## Abstract

Plant-derived nanovesicles (PDNVs) have attracted increasing attention for their potential biomedical applications, particularly in oxidative stress-related diseases and neuroprotection. *Wolffia globosa*, an emerging functional food and potential future food source, is rich in phytochemicals associated with health-promoting properties. Harnessing *W. globosa*–derived nanovesicles (WGNVs) represents a promising strategy for expanding the biomedical applications of this plant. In this study, WGNVs were isolated and characterized, and their neuroprotective potential was evaluated in vitro. The isolated WGNVs exhibited nanoscale dimensions and a lipid bilayer structure. Biochemical analyses revealed substantial total phenolic and flavonoid contents, alongside intrinsic antioxidant activities. Untargeted metabolomic profiling identified a diverse repertoire of metabolites, including major constituents subsequently validated using independent analytical approaches. These findings provided a foundation for investigating the neuroprotective effects of WGNVs in a methylglyoxal (MGX)-induced model of neuronal oxidative stress. High-resolution confocal microscopy demonstrated the cellular localization of WGNVs within neuronal cells, providing evidence of interactions between plant-derived nanovesicles and neuronal cells. WGNV treatment attenuated MGX-induced free radical accumulation and oxidative stress responses while preserving cellular morphology integrity. Collectively, this study provides foundational evidence of the physicochemical characteristics, metabolite composition, and in vitro neuroprotective potential of WGNVs. These findings highlight WGNVs as a promising naturally derived nanocarrier platform for investigating plant–animal interkingdom interactions and exploring potential therapeutic applications across a broad range of biomedical fields.

**Highlights:**

- *Wolffia globosa*–derived nanovesicles (WGNVs) exhibited nanoscale sizes, lipid bilayer architecture, and intrinsic antioxidant activities.
- LC–MS-based metabolomics revealed diverse chemical signatures spanning 227 metabolite classes, including fatty acid derivatives, dipeptides, flavonoid 8-C-glycosides, and triterpenoids.
- WGNVs were internalized by mouse neuronal CAD cells, highlighting their biocompatibility and potential to facilitate cross-kingdom communication.
- WGNVs alleviated methylglyoxal-induced oxidative stress and maintained neuronal morphological integrity in the neuronal cells, highlighting their potential neuroprotective relevance to diabetes- and aging-associated neurodegeneration.
- This study highlights *W. globosa* as a promising source of bioactive nanovesicles, bridging future food research and nanomedicine development.

## 1. Introduction

Functional foods have attracted increasing scientific interest not only for their nutritional benefits but also for their potential as valuable sources of naturally occurring bioactive materials for biomedical applications. In particular, exploring innovative approaches to harnessing these biological resources offers promising opportunities to extend their value beyond conventional nutritional applications. In Thailand, this approach aligns with the Bio-Circular-Green (BCG) Economy model, which promotes the sustainable utilization of biological resources to develop high-value products through scientific and technological innovation.

Within this context, *Wolffia globosa*, commonly known as duckweed, watermeal, or Khai Pham in Thailand, is among the smallest flowering aquatic plants in the Lemnaceae family and has attracted increasing attention as a promising future functional food owing to its rich nutritional composition, rapid growth, and potential for sustainable cultivation[1–4]. *W. globosa* is recognized for its protein content, essential amino acids, vitamins, minerals, and diverse phytochemical constituents. Beyond its nutritional value, *W. globosa* contains naturally occurring secondary metabolites, particularly phenolic compounds and flavonoids, associated with antioxidant properties[1]. These characteristics highlight the potential of *W. globosa* as a valuable biological resource for developing novel bioactive materials with biomedical applications. However, previous studies have primarily focused on its nutritional composition and phytochemical properties, whereas the biological potential of its naturally occurring nanovesicles remains largely unexplored.

Plant-derived nanovesicles (PDNVs) are plant-origin nanoscale vesicles that share structural and biochemical features with mammalian exosomes and have gained increasing attention for their potential biomedical applications [5]. These vesicular structures are generally characterized by lipid bilayer membranes and associated with diverse molecular constituents, including nucleic acids, proteins, and secondary metabolites. Their membrane-associated organization enables them to function as natural bioorganic excipients, providing a biocompatible platform that facilitates interkingdom interactions between plant-derived molecular components and recipient mammalian cells[6]. Recent investigations have demonstrated that nanovesicle preparations isolated from edible and medicinal plants exhibit various biological activities, including antioxidant, anti-inflammatory, anti-cancer, and cytoprotective effects[7–11]. However, the physicochemical properties, molecular composition, and biological activities of PDNVs can vary considerably depending on their botanical origin and isolation procedures. Therefore, the isolation and characterization of nanovesicles from emerging plant sources, such as *W. globosa*, are essential for elucidating their structural properties, molecular composition, and potential biological functions.

The biological activities of PDNVs are influenced by their diverse molecular constituents, particularly phenolic compounds and flavonoids with potential antioxidant and cellular regulatory properties. However, conventional targeted analyses may overlook the broader chemical diversity associated with these nanovesicles. An unbiased, untargeted metabolomic approach using liquid chromatography–quadrupole time-of-flight tandem mass spectrometry (LC–QTOF–MS/MS) enables comprehensive profiling and annotation of bioactive metabolites. Previous LC–QTOF–MS/MS-based phytochemical profiling of *W. globosa* biomass has revealed diverse phenolic constituents associated with antioxidant properties[2,3,12]. Nevertheless, previous investigations have primarily focused on whole-plant extracts, whereas the metabolomic composition of *W. globosa*-derived nanovesicle (WGNVs) and its potential relevance to their biological activities remain largely unexplored.

Recent studies have established a foundation for WGNV research, with one reporting their successful isolation and characterization[13] and another demonstrating their potential anticancer activity against breast cancer cells[14]. Despite their emerging biomedical potential, the metabolite composition of WGNVs remains uncharacterized, and their potential cytoprotective properties have yet to be investigated. Oxidative stress is a major contributor to cellular dysfunction and has been implicated in the development and progression of aging-associated and neurodegenerative disorders. Excessive accumulation of reactive oxygen species (ROS) can disrupt cellular redox homeostasis, alter stress-responsive gene expression, and compromise cellular structural integrity. Methylglyoxal (MGX), a highly reactive dicarbonyl metabolite primarily generated as a byproduct of glycolysis, can accumulate under pathological conditions associated with aging, diabetes, and neurodegenerative disorders[15,16]. Elevated MGX levels promote the formation of advanced glycation end products (AGEs), mitochondrial dysfunction, and excessive ROS generation, thereby exacerbating oxidative stress and cellular damage. Furthermore, MGX exposure has been associated with neurotoxicity, potentially contributing to neurodegenerative processes[17–21]. Consequently, MGX-induced neuronal stress represents a relevant experimental model for screening bioactive compounds with neuroprotective potential, as demonstrated by previous investigations of naringenin[22], sulforaphane[23] and tenuigenin[24], and provides a foundation for identifying promising candidates for biomedical applications targeting oxidative stress-associated neurological disorders.

In the present study, we report the isolation and characterization of WGNVs and investigated their molecular composition and neuroprotective potential in vitro. An unbiased, untargeted LC–QTOF–MS/MS-based metabolomic approach was employed to identify the bioactive metabolites of WGNVs. Furthermore, their neuroprotective properties were evaluated in a neuronal cell model of MGX-induced oxidative stress. Beyond its conventional nutritional applications, this study highlights the potential of harnessing *W. globosa*–derived nanovesicles as naturally occurring bioactive platforms, promoting the value-added utilization of this emerging future functional food and opening new avenues for biomedical innovation.

## 2. Material and methods

### 2.1 Cell culture

The mouse CNS-derived CAD cell line was obtained from the European Collection of Authenticated Cell Cultures (ECACC, accession number: 08100805) and cultured in Dulbecco’s modified Eagle’s medium (DMEM): nutrient mixture F-12 (F12) (Gibco, USA) supplemented with 8% fetal bovine serum (FBS) (HyClone, USA) and 1% antibiotic-antimycotic 100X (Gibco, USA)[25,26]. The human neuroblastoma SH-SY5Y (cat.no. iCell-h187) cell line was purchased from iCell Bioscience, China. The cells were maintained in the DMEM: F12 supplemented with 10% (v/v) FBS and 1% (v/v) antibiotic-antimycotic 100X[27]. The cells were maintained in a humidified incubator at 37 °C with 5% CO2 and 95% air (v/v). The cultured cells were regularly tested for mycoplasma contamination via a Mycoplasma PCR detection kit (Abcam, UK).

### 2.2 Isolation of *Wolffia globosa*–derived nanovesicles through a combined ultracentrifugation and ultrafiltration

Fresh, cleaned whole plants of *W. globosa* were purchased from a local plant market that employed a controlled, high-quality cultivation system. The plant material was identified by Dr. Panitch Boonsnongcheep. Nanovesicles were isolated using a sequential centrifugation procedure adapted from our previously established protocol [27]. The diagram is depicted in Figure 1c. Briefly, approximately 2 kg of plant material was mixed with sterile 1x phosphate-buffered saline (PBS) at a 1:1 (w/v) ratio and homogenized using a blender. The resulting homogenate was sequentially filtered through white filter paper to remove large fibrous materials. The filtrate was then centrifuged at 10,000 *x g* for 60 min at 4 °C for two consecutive rounds to remove residual cellular debris. The resulting supernatants were collected and further centrifuged at 50,000 *x g* for 60 min at 4 °C for two consecutive rounds. The supernatants were subsequently passed through a 0.45-μm filter and subjected to ultracentrifugation at 100,000 *x g* for 60 min at 4 °C for two consecutive rounds to pellet the nanovesicles. To reduce potential contaminants, the resulting pellets were subjected to ultrafiltration using Macrosep centrifugal filters (MAP100C37, Cytiva, USA) to remove low-molecular-weight components and soluble contaminants. The retained fraction was subsequently centrifuged at 3,000 *x g* for 15 min for three consecutive rounds. The final suspension, with a volume of approximately 10 mL, was passed through a 0.22-μm membrane filter and stored at −80 °C until further use.

**Figure 1.**
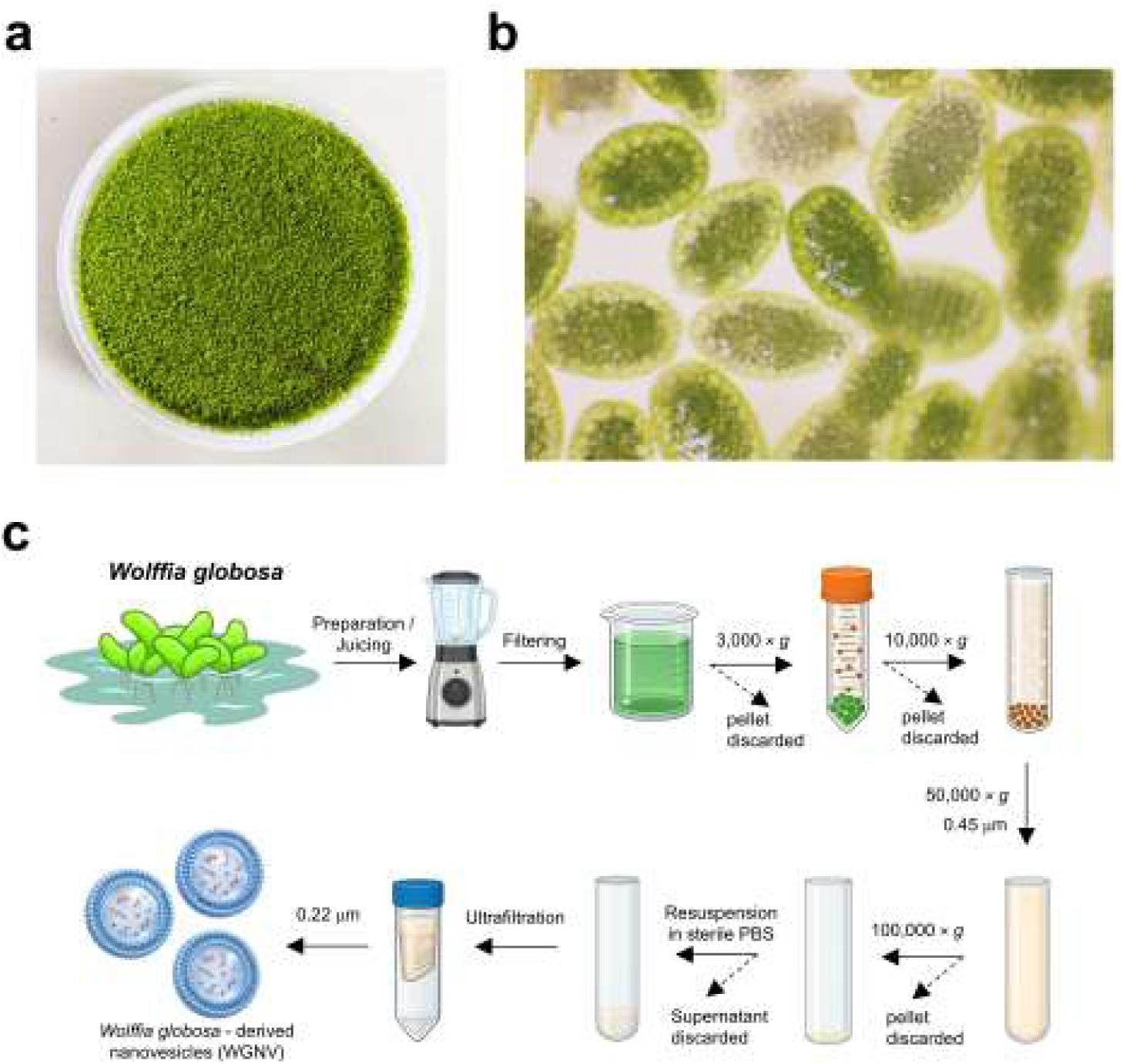
Morphology of *Wolffia globosa* and the isolation procedure for *W. globosa*–derived nanovesicles (WGNVs). **(a)** Macroscopic appearance of fresh *W. globosa*. **(b)** Microscopic appearance of individual *W. globosa* plants, showing their characteristic globular morphology (5X objective lens). **(c)** Schematic illustration of the WGNV isolation procedure involving plant preparation and juicing, filtration, sequential centrifugation. The resulting pellet was resuspended and subjected to ultrafiltration for WGNVs purification.

### 2.3 Nano-tracking particle analysis (NTA)

The size distribution and concentration of the WGNVs were analyzed via NanoSight Pro (Malvern Panalytical, UK). The samples (200 μL) were injected with a sterile syringe. For each measurement, three to five videos were captured. The measurements were performed at room temperature.

### 2.4 Zeta potential analysis

Zeta potential analysis was performed to determine the surface charge and colloidal stability of the isolated nanovesicles[28]. The sample was diluted in sterile distilled water (1 mL) and applied to the nanoPartica SZ-100V2 nanoparticle analyzer (HORIBA, Japan). Zeta potential analysis was carried out using standard setting (dispersion medium viscosity = 0.896 mPas, temperature = 25 °C). The data were analyzed by the Horiba SZ-100 software version 2.44.

### 2.5 Cryogenic electron microscope (Cry-EM)

The samples containing the isolated *W. globosa*-derived nanovesicles were submitted to the Research Frontier Facility (FRF), Mahidol University, for structural analysis. An aliquot of each sample (3 μL) was applied to a plasma-cleaned copper grid (C-flat R 1.2/1.3, 300 mesh, Electron Microscopy Sciences) and prepared via an automated grid plunger (Vitrobot Mark IV, Thermo Fisher Scientific) with the environmental chamber set at 100% humidity and 4 °C. The grid was blotted for 4.5 s and vitrified in liquid ethane. The frozen hydrated WGNVs were imaged via a cryogenic electron microscope (Glacios, Thermo Fisher Scientific) equipped with a Falcon 3EC. Cryo-EM imaging was performed at 73,000x and 120,000x magnification at 200 kV.

### 2.6 DPPH radical scavenging assay

The antioxidant activity was monitored by the reduction reaction of the DPPH free radical (2,2-diphenyl-1-picrylhydrazyl). The samples were prepared with a 2-fold dilution from the highest concentration of WGNVs. The sample (100 μL) was mixed with 100 μL of DPPH solution (0.2 mM) freshly dissolved in methanol and incubated at room temperature in the dark for 30 min. Trolox (Sigma Aldrich, USA) was used as a standard antioxidant. The color change of the solution from an intense purple color to pale yellow was observed, and the absorbance (A) values were detected at 517 nm via a SpectraMax iD5e multi-mode microplate reader and the SoftMax Pro data acquisition and analysis software version 7.4 (Molecular Devices, USA). The DPPH radical-scavenging activity was calculated using the formula described in our previous work[27]. The Trolox equivalent (μg/mL) concentrations were obtained from the linear regression equation of the Trolox calibration curve.

### 2.7 Ferric reducing antioxidant power (FRAP)

The ability of WGNVs to donate electrons and reduce ferric iron (Fe^3+^) to ferrous iron (Fe^2+^) at low pH was evaluated [29]. FRAP reagent was freshly prepared by mixing 100 mM acetate buffer pH 3.6, 10 mM 2,4,6-Tris(2-pyridyl)-striazine in 40 mM HCl, and 20 mM FeCl3 (10:1:1). The samples and standards were prepared in the similar manners as in the DPPH assay. The reaction mixture composed of 20 μL of samples or standards and 180 μL of reaction mixture. The absorbance was measure at 595 nm using the same equipment as mention previously. The ferric reducing antioxidant power (FRAP) was determined based on absorbance measurements and expressed as Trolox equivalents (µg/mL).

### 2.8 Total flavonoid content (TFC) measurement

The aluminum chloride assay was used for determination of total flavonoid contents[30]. Quercetin was used as the standard flavonoid. It was prepared at the concentration of 1.56 – 100.00 μg/mL. Samples were prepared as mentioned in the antioxidant assays. In each well of a 96-well plate, 50 μL of standard or samples were mixed with 150 μL of 10% aluminum chloride. After incubation for 6 min at room temperature, the absorbances were measured at 430 nm. A standard curve was constructed by plotting the measured absorbance values against the corresponding quercetin concentrations. The contents of flavonoids in samples were reported as μg/mL quercetin equivalent.

### 2.9 Total phenolic content (TPC) measurement

Total phenolic content of WGNVs were measured according to Folin-Ciocalteu assay[31,32]. Gallic acid was used as the standard phenolic compound and prepared at the concentration of 1.56 – 100.00 µg/mL. Samples were prepared as previously mentioned. 20 _μ_L of standards or samples were mix with 100 µL of Folin-Ciocalteu reagent, and 80 _μ_L 7.5% sodium carbonate, and after incubation at room temperature for 30 min, absorbances were measured at 760 nm. Standard curve was plotted using concentrations of gallic acid and absorbances, and the levels of phenolic compounds in WGNVs were reported as µg/mL gallic acid equivalent.

### 2.10 Acetylcholinesterase activity assay

Human SH-SY5Y cells were seeded in 60-mm culture dishes at a density of 0.3 × 10^6^ cells per dish and treated with WGNVs (10^7^ particles/mL) for 24 h. Acetylcholinesterase (AChE) activity was assessed according to a previously published protocol[33]. Following treatment, the cells were harvested, and total cell lysates were homogenate in 1x PBS containing 1% Triton X-100. Protein concentrations were determined using a Pierce BCA protein assay kit (cat no. 23225; Thermo Scientific, USA), and 20 μg of total protein per sample was used to measure AChE activity using the commercial kit (MAK119, Sigma-Aldrich, USA), according to the manufacturer’s instructions. Absorbance was measured at 412 nm using a SpectraMax iD5e multi-mode microplate reader. AChE activity was calculated relative to that of the untreated control. The experimental workflow is illustrated in Supplementary Figure 1.

### 2.11 Untargeted metabolomic analysis

Metabolite profiling of lyophilized *W. globosa*–derived nanovesicles was performed using a liquid chromatography–quadrupole time-of-flight mass spectrometry (LC– QTOF–MS) platform, following a previously published protocol[27]. Briefly, samples were extracted with 200 µL of 70% methanol containing sulfadimethoxine (50 ng/mL) as an internal standard to enable subsequent signal normalization. The mixture was thoroughly vortexed and subjected to ultrasonic extraction for 15 min at room temperature to facilitate metabolite solubilization. Following extraction, the samples were centrifuged at 14,000 rpm for 10 min, and the clarified supernatants were carefully transferred into liquid chromatography–mass spectrometry (LC–MS) vials for instrumental analysis.

Chromatographic separation was conducted on an Agilent LC system equipped with a Poroshell 120 EC-C18 column (2.1 × 100 mm, 2.7 µm particle size) maintained at 50°C. The injection volume was set to 10 µL, and the flow rate was maintained at 0.4 mL/min throughout the analysis. Mass spectrometric detection was performed via an Agilent LC-QTOF 6545XT operated in high-resolution acquisition mode under both positive and negative electrospray ionization (ESI) conditions. The raw data were processed via MS-DIAL software version 5.3. Metabolite annotation considered M+H adducts in positive mode and M−H adducts in negative mode. Metabolite identification was carried out by matching accurate mass, retention time, and MS/MS fragmentation patterns against multiple spectral libraries, including the MS-DIAL ESI (+/−) MS/MS database from authentic standards [34], the Fiehn/Vaniya natural product library, and the BMDMS-NP database [35] The identification criteria included a retention time window of 0.2–18 min, a sample-to-blank ratio of at least 100, an identification score ≥ 0.7, and a mass error tolerance within ± 25 ppm. Using these stringent criteria, a total of 280 metabolic features were detected in positive ionization mode, and 184 features were detected in negative ionization mode. Detailed information on the annotated metabolites is provided in Supplementary File 1.

### 2.12 HPLC analysis

All reference standards including Schaftoside, isoorientin, orientin, and isovitexin were purchased from Biopurify Phytochemicals (Chengdu, China). The selected metabolites were validated using a Thermo Scientific UltiMate 3000 high-performance liquid chromatography (HPLC) system equipped with a diode-array detector (DAD). Isocratic elution was performed using a mobile phase consisting of 20% acetonitrile acidified with 2.5% acetic acid at a flow rate of 0.5 mL/min. Chromatographic separation was achieved using a Cosmosil C18 reversed-phase column (4.6 × 150 mm; Nacalai Tesque, Japan). Chromatograms were recorded at 254 nm.

### 2.13 Cell viability

The CAD cells were plated on a 96-well plate at a density of 5,000 cells/well. Then, the cells were exposed to different concentrations of the WGNV (particle/mL) or methylglyoxal (MGX) solution (M0252, Sigma-Aldrich, USA) for 24 and 48 h, as indicated. A control group was included in which cells received the same final volume of PBS or water used to dilute the nanovesicles and MGX, respectively. At the end of the experiment, cell viability was determined via an MTT assay (M5655, Sigma Aldrich, USA). The MTT stock reagent (5 mg/mL) was directly added to the conditioning media to obtain the final concentration (0.5 mg/mL) of the MTT reagent. The plates were further cultured for an additional 3 h. The MTT solution was carefully removed, and insoluble formazan was dissolved in DMSO. Colorimetric determination was performed at 570 nm via a SpectraMax iD5e multi-mode microplate reader. The data were normalized to the control data and are presented as the relative absorbance.

### 2.14 The cellular uptake assay

The PKH26 red fluorescent cell linker mini kit for general cell membrane labeling (MINI26, Sigma Aldrich, USA) was used to observe the cellular localization of WGNV by neuronal cells. The mixture was combined with 400 _μ_L of the nanovesicles, 8 _μ_l of PKH26 dye, and 0.5 mL of Diluent C and incubated for 5 min at room temperature. The reaction was terminated by adding 10 mL of 1% bovine serum albumin (BSA). The collection of labeled particles and removal of excess free dye were performed via Macrosep advance centrifugal devices as described previously. Then, the concentrated PHK26-tagged WGNV (~1.5 ml) were collected, resuspended in complete culture media, and incubated with the neuronal cells overnight. The cells were fixed, permeabilized, and stained for βIII-tubulin and DAPI as described in the immunocytochemistry section.

### 2.15 Immunocytochemistry

The cells were fixed with 4% paraformaldehyde (PFA) in 1x PBS for 20 min at room temperature and rinsed three times with 1x PBS. The cells were then permeabilized in 0.2% Triton X-100 for 15 min at room temperature. The plates were washed with PBS three times. Nonspecific binding was blocked with 1% BSA in PBS for 30 min at 37 °C. The cells were stained with anti-βIII-tubulin (T8578, Sigma Aldrich, USA) on parafilm for 30 min at 37 °C before being washed with PBS three times, each for more than 5 min. The cells were subsequently incubated with donkey anti-mouse IgG-FITC (cat.no. sc-209, Santa Cruz Biotechnology, USA) for 30 min at 37 °C. The final dilution was 1:400 for all the antibodies. The incubated cells were washed with PBS three times, washed with clean water, and mounted on slides using Fluoromount-G Mounting Medium with DAPI (cat. no. 00495952; Invitrogen, USA). Images were acquired via a Zeiss LSM 800 confocal laser scanning microscope.

For cell shape analysis, images were acquired from multiple non-overlapping fields and processed using ImageJ software version 1.54g (National Institutes of Health, USA). Cell area (μm^2^), cell roundness, and solidity were quantified according to a previously published method[36].

### 2.16 Measurement of intracellular reactive oxygen species (ROS)

Intracellular ROS levels were assessed using 2′,7′-dichlorodihydrofluorescein diacetate (DCFH-DA), following a previously published protocol [37]. Neuronal CAD cells were seeded in 96-well plates at a density of 5,000 cells per well and cultured for 24 h. The cells were then exposed to 200 _μ_L of treatment medium containing WGNVs or MGX at the indicated concentrations, as specified in the figure legends. Following the treatment period, the medium was removed, and 200 _μ_L of fresh culture medium containing 10 _μ_M DCFH-DA (D6883, Sigma-Aldrich, USA) was added to each well. The plate was incubated at 37 °C for 30 min in the dark. The DCFH-DA working solution was then removed, and the cells were washed once with 1XPBS.

For quantitative analysis, 100 μL of RIPA buffer (R0278, Sigma-Aldrich, USA) was added to each well. After incubation on ice for 5 min, the cell lysates were collected into 1.5-mL microcentrifuge tubes and centrifuged at 21,130 x g for 10 min at 4 °C. The resulting supernatants were transferred to a black 96-well plate, and fluorescence intensity was measured using a SpectraMax iD5e multimode microplate reader (excitation/emission wavelengths: 485/530 nm). Fluorescence intensity was normalized to the total protein concentration, determined using the BCA assay. For fluorescence imaging, the DCFH-DA stained cells were washed with 1X PBS and imaged using a Zeiss LSM 800 confocal laser-scanning microscope.

### 2.17 Quantitative real-time polymerase chain reaction (qRT PCR)

Total RNA was extracted with PureLink RNA Mini Kit (cat no. 12183018A, Invitrogen, Carlsbad, USA) according to the manufacturer’s instructions. The concentration and purity of the extracted RNA were determined via a NanoDrop One Spectrophotometer (Thermo Fisher Scientific, USA). cDNA synthesis was conducted via an iScript cDNA Synthesis Kit (Bio-Rad Laboratories, USA). Total cDNA (200 ng in 2 μl) was mixed with 8 μl of iTaq Universal SYBR Green Supermix (Bio-Rad Laboratories, USA) containing specific forward and reverse primers (400 nM each) to perform real-time PCR amplification, following the manufacturer’s instructions. The relative mRNA expression of the target genes was normalized to that of GAPDH as the internal control and was calculated via the 2^−ΔΔCt^ method. A list of primers and their sequences for targeting genes in mice (Mus Muscularis) is provided in Table 1.

**Table 1.**
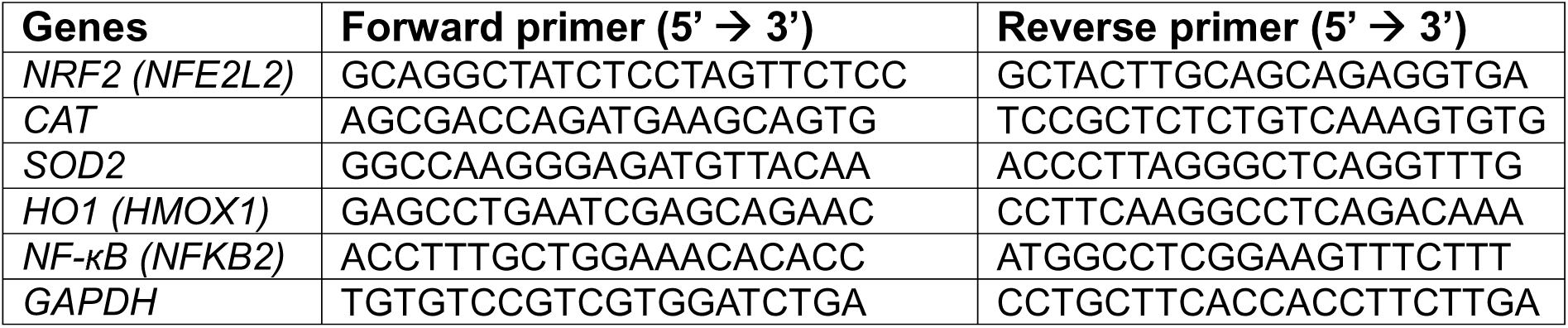
Primer sequences targeting genes in *Mus musculus* used in this study.

### 2.18 Statical analysis

Statistical analysis was performed, and graphs were created via GraphPad Prism version 8.0.1. Statistical significance was determined via one-way ANOVA for multiple comparisons. Statistical significance was interpreted at a p value < 0.05. All experiments were performed with at least three biological replicates.

## 3. Result

### 3.1 *Wolffia globosa–*derived nanovesicles exhibited a nanoscale size distribution and a lipid bilayer structure

*W. globosa* is one of the smallest flowering aquatic plants, exhibiting a globular shape (Figure 1a–b). Freshly harvested and cleaned *W. globosa* was subjected to nanovesicle isolation using a combination of high-speed centrifugation and ultrafiltration, as illustrated in Figure 1c.

To confirm successful nanovesicle isolation, the *W. globosa*-derived nanovesicles (WGNVs) were characterized using nanoparticle tracking analysis (NTA) to determine particle size distribution and concentration, zeta potential measurement to assess surface charge, and cryogenic transmission electron microscopy (cryo-EM) to examine structural characteristics. NTA revealed that the isolated particles were distributed within the nanometer range, with mode and mean values of 127.5 nm and 187 nm, respectively (Figure 2a-b). The results showed that the D50 was 154 nm, indicating that 50% of the measured particles had diameters of 154 nm or smaller. Whereas only 10% had diameters greater than 291 nm (D90). The unimodal particle size distribution suggested that the isolated WGNVs exhibited a relatively homogeneous size profile. The isolation procedure yielded a high particle concentration of up to 10^11^ particles/mL. The zeta potential of the isolated WGNVs was −24.7 mV, indicating a negatively charged particle surface and suggesting moderate electrostatic repulsion between particles, which may contribute to colloidal stability (Figure 2c). The clearly visible bilayer membrane structure in the cryo-EM images (Figure 2d) supports the vesicular morphology of the isolated WGNVs. Overall, the plant-derived nanovesicles were successfully isolated and characterized from *W. globosa*, supporting their suitability for subsequent downstream experiments.

**Figure 2.**
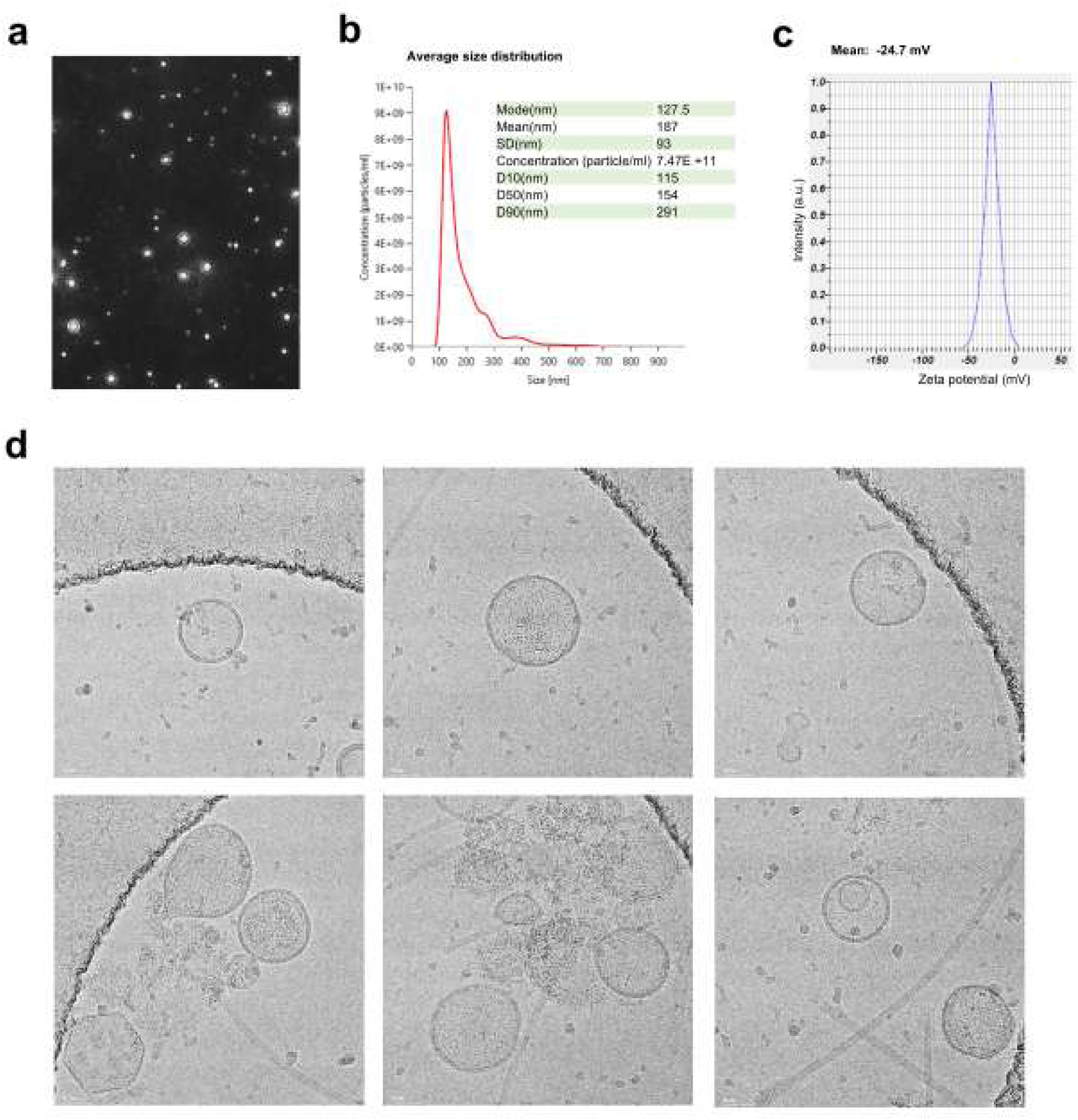
Characterization of *W. globosa*–derived nanovesicles revealed nanoscale particles enclosed by a bilayer membrane. **(a)** Representative nanoparticle tracking analysis (NTA) video frames displaying real-time light scattering during particle Brownian motion. The full video is available in Supplementary file 1. **(b)** Particle size distribution and related parameters measured by NTA. **(c)** Zeta potential measurement of WGNVs. **(d)** Representative cryo-EM images showing the vesicular morphology and bilayer membrane structure of WGNVs. Scale bar is 50 μm.

### 3.2 WGNVs possessed intrinsic antioxidant properties

To screen the bioactivities of the isolated WGNVs, we evaluated their intrinsic free radical scavenging capability via DPPH and FRAP, two common assays for measuring the antioxidant potential. Trolox, a cell-permeable analog of vitamin E with potent antioxidant properties was used as the reference standard (Figure 3).

**Figure 3.**
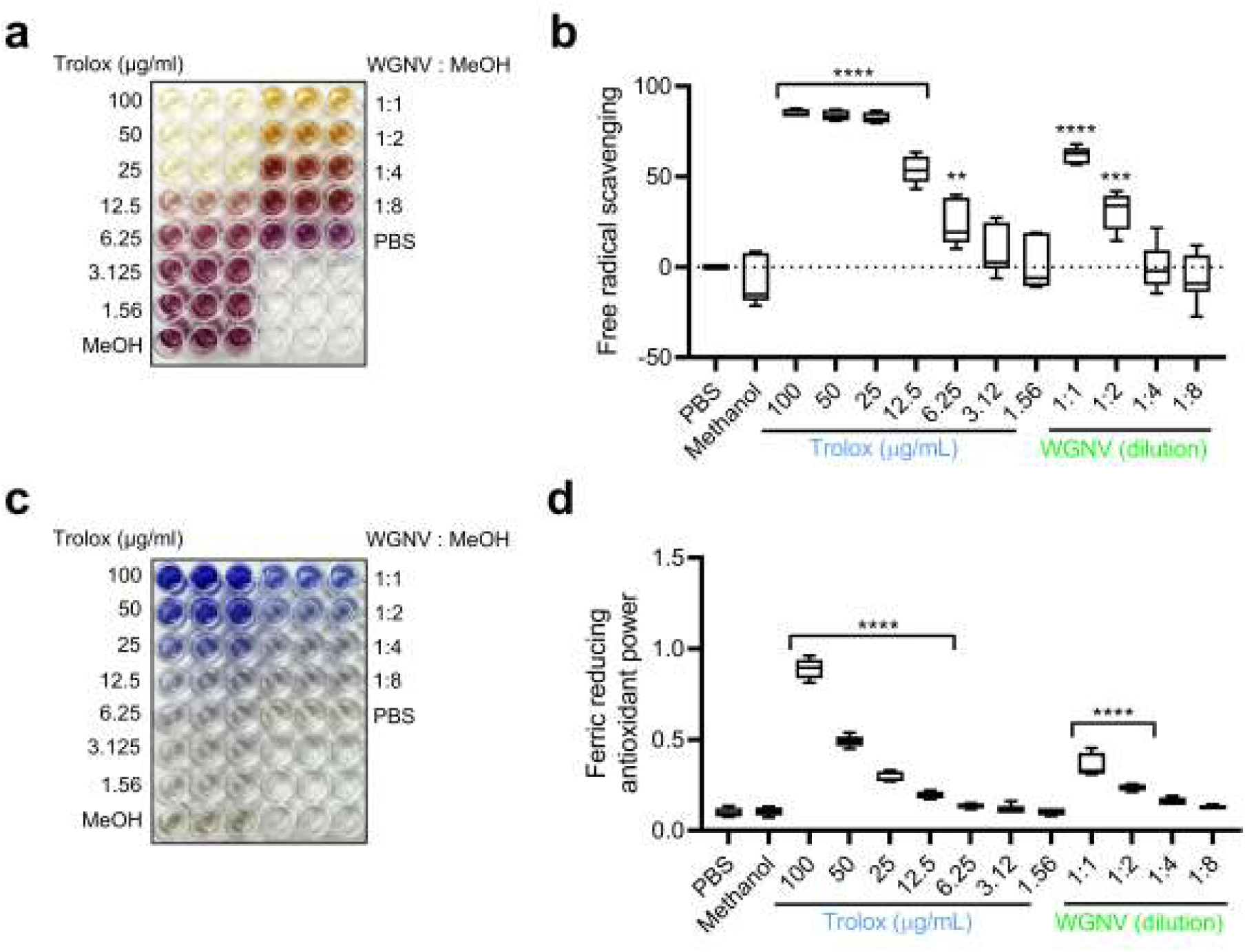
Intrinsic antioxidant properties of WGNVs in cell-free assays. **(a)** Representative images of the DPPH assay showing color changes that indicate the free radical scavenging activity of Trolox (standard antioxidant) and WGNVs at different dilution ratios in methanol. **(b)** Quantification of DPPH free radical scavenging activity, expressed as a percentage relative to the PBS control. **(c)** Representative images of the FRAP assay showing the formation of an intense blue ferrous–tripyridyltriazine (Fe^2+^– TPTZ) complex, indicating the reducing capacity of Trolox (standard antioxidant) and WGNVs at different dilution ratios in methanol. **(d)** Quantification of ferric reducing antioxidant power (FRAP) based on optical density measured at 595 nm. Data are presented as mean ± SD from six replicates (n = 6). Statistical significance was determined using one-way ANOVA followed by Tukey’s multiple comparisons test. **p < 0.01, ***p < 0.001, and ****p < 0.0001 (vs the PBS control).

The DPPH assay measures free radical scavenging activity through the colorimetric transition of the DPPH solution from deep purple to pale yellow. As shown in Figure 3a, increasing concentrations of Trolox (1.56-100 μg/mL) and higher concentrations of WGNVs (1:1 to 1:8 dilutions in methanol) exhibited a dose-dependent reduction in purple color intensity. Quantitative analysis (Figure 3b) showed that Trolox significantly scavenged free radicals, reaching maximum scavenging efficiency (up to 90%) at 25-100 μg/mL. Similarly, incubation with WGNVs conferred significant radical scavenging capacity in a dose-dependent fashion. The 1:1 diluted concentration achieved 62 % free radical scavenging activity, while the 1:2 dilution maintained statistically significant activity.

Furthermore, the reducing capacity of the test samples was evaluated using the FRAP assay by monitoring the formation of the intense blue tripyridyltriazine complex. Higher concentrations of both Trolox and WGNVs produced stronger blue color intensity on the assay plate (Figure 3c). Quantitative readout (Figure 4d) confirmed that Trolox exhibited a robust, concentration-dependent increase in reducing potential, peaking at an absorbance value near 0.9 at 100 μg/mL. In parallel, WGNVs displayed a significant, concentration-dependent ferric reducing capacity. The 1:1 and 1:2 WGNV preparations yielded significantly higher reducing power compared to control vehicles, while lower concentrations showed baseline values.

**Figure 4.**
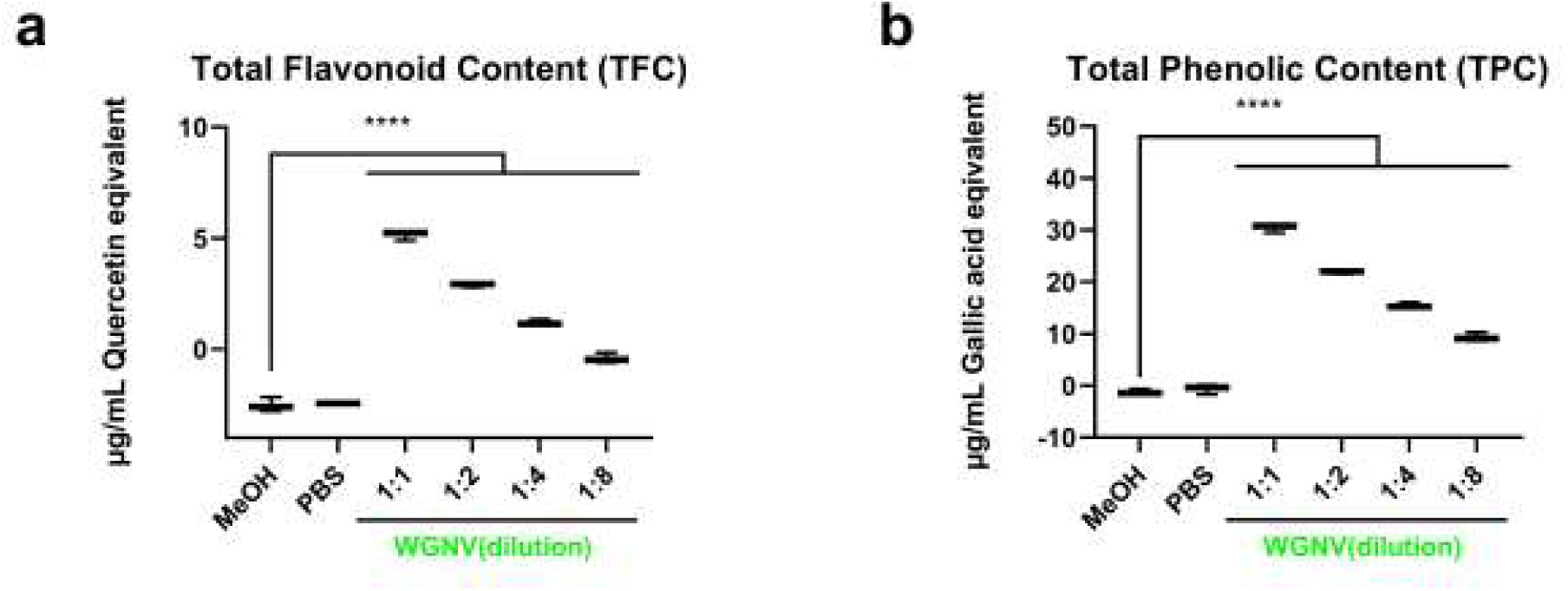
The *W. globosa*–derived nanovesicles contained measurable levels of total flavonoids and phenolic contents. **(a)** Total flavonoid content (TFC), expressed as μg/mL quercetin equivalents. **(b)** Total phenolic content (TPC), expressed as μg/mL gallic acid equivalents. WGNV samples were evaluated at different dilution ratios (1:1, 1:2, 1:4, and 1:8), with methanol (MeOH) and PBS included as controls. Data are presented as mean ± SD from three replicates (n=3) Statistical significance was determined using one-way ANOVA followed by Tukey’s multiple comparisons test. ****p < 0.0001 (vs the control).

Table 2 summarizes the Trolox-equivalent antioxidant capacity of WGNVs determined using two assays. At a 1:1 concentration, WGNVs exhibited antioxidant capacities equivalent to 17.67 μg/mL and 30.52 μg/mL of Trolox, as determined by the DPPH and FRAP assays, respectively. Taken together, these findings demonstrate that WGNVs possess intrinsic antioxidant activity, providing a foundation for further metabolite profiling to identify potentially bioactive compounds and for evaluating their protective effects in cellular models of oxidative stress.

**Table 2.** Trolox-equivalent antioxidant capacity of WGNVs determined using DPPH and FRAP assays. The Trolox-equivalent concentrations (μg/mL) were calculated using the linear regression equations derived from the respective Trolox calibration curves. Data are presented as mean ± standard deviation (SD).

| **WGNV (ratio)** | **Trolox eq (µg/mL)** |  |
|---|---|---|
|  | **DPPH** | **FRAP** |
| 1:1 | 17.67±0.86 | 30.52±2.17 |
| 1:2 | 11.54±7.56 | 20.44±2.93 |

### 3.3 *W. globosa*–derived nanovesicles represent a potential natural source of bioactive phenolic and flavonoid compounds

The antioxidant activity of the isolated WGNVs suggests the presence of bioactive metabolites with potential antioxidant properties. Phenolic compounds and flavonoids are important classes of phytochemicals that contribute to antioxidant activity. Therefore, the total phenolic content (TPC) and total flavonoid content (TFC) of the isolated WGNVs were subsequently assessed.

The TFC was expressed as μg/mL quercetin equivalents (Figure 4a). The 1:1 diluted WGNV exhibited the highest flavonoid concentration at approximately 5 μg/mL quercetin equivalents, with levels decreasing in a clear dose-dependent manner to roughly 3 μg/mL (1:2) and 1 μg/mL (1:4). In parallel, Figure 4b presents the TPC expressed as μg/mL Gallic acid equivalents. Consistent with the flavonoid profile, WGNVs demonstrated a strong, concentration-dependent abundance of total phenolic compounds. The highest concentration was detected in the 1:1 dilution (~30 μg/mL Gallic acid equivalent), followed by a step-wise decline across 1:2 (~20 μg/mL Gallic acid equivalent), 1:4 (~15 μg/mL Gallic acid equivalent), and 1:8 (~9 μg/mL Gallic acid equivalent) dilutions. The TPC values across all WGNV preparations were significantly higher than those observed in the MeOH and PBS negative controls.

These findings indicate that WGNVs contain phenolic and flavonoid compounds. The presence of these phytochemicals may explain the antioxidant activity observed in the DPPH and FRAP assays, highlighting the potential of WGNVs as cargo carriers of naturally occurring antioxidant compounds.

### 3.4 Untargeted LC-QTOF-MS/MS based metabolomic analyze identified the composition of bioactive molecules in *W. globosa*-derived nanovesicles

To identify the potential bioactive metabolites in WGNVs, untargeted LC-QTOF-MS/MS-based metabolomic approach was performed. Chromatogram profile obtained from positive and negative ionization modes is shown in Supplementary Figure 1. A total of 464 metabolites were identified and classified into 227 ontology classes (Supplementary File 2). The data highlight the chemical diversity of WGNVs and their potential as a rich source of naturally occurring bioactive molecules.

The top 25 metabolite ontology classes are presented in Figure 5. Six classes contained more than 10 identified metabolites, including linoleic acids and derivatives, dipeptides, long-chain fatty acids, flavonoid 8-C-glycosides, medium-chain fatty acids, and triterpenoids. Notably, the identification of flavonoid 8-C-glycosides, together with other flavonoid-related classes, including phenolic glycosides, flavonoid C-glycosides, flavonoid 3-O-glycosides, and flavonoid 7-O-glycosides, highlights the diversity of flavonoid-related metabolites in WGNVs. Representative compounds included orientin isoorientin, vitexin, isovitexin, schaftoside, and isoschaftoside as well as the flavonoids quercetin and kaempferol. These findings are consistent with the high total flavonoid and phenolic content measured in WGNVs, supporting their potential as a natural source of bioactive phytochemicals that may contribute to the antioxidant activity observed in the DPPH and FRAP assays.

**Figure 5.**
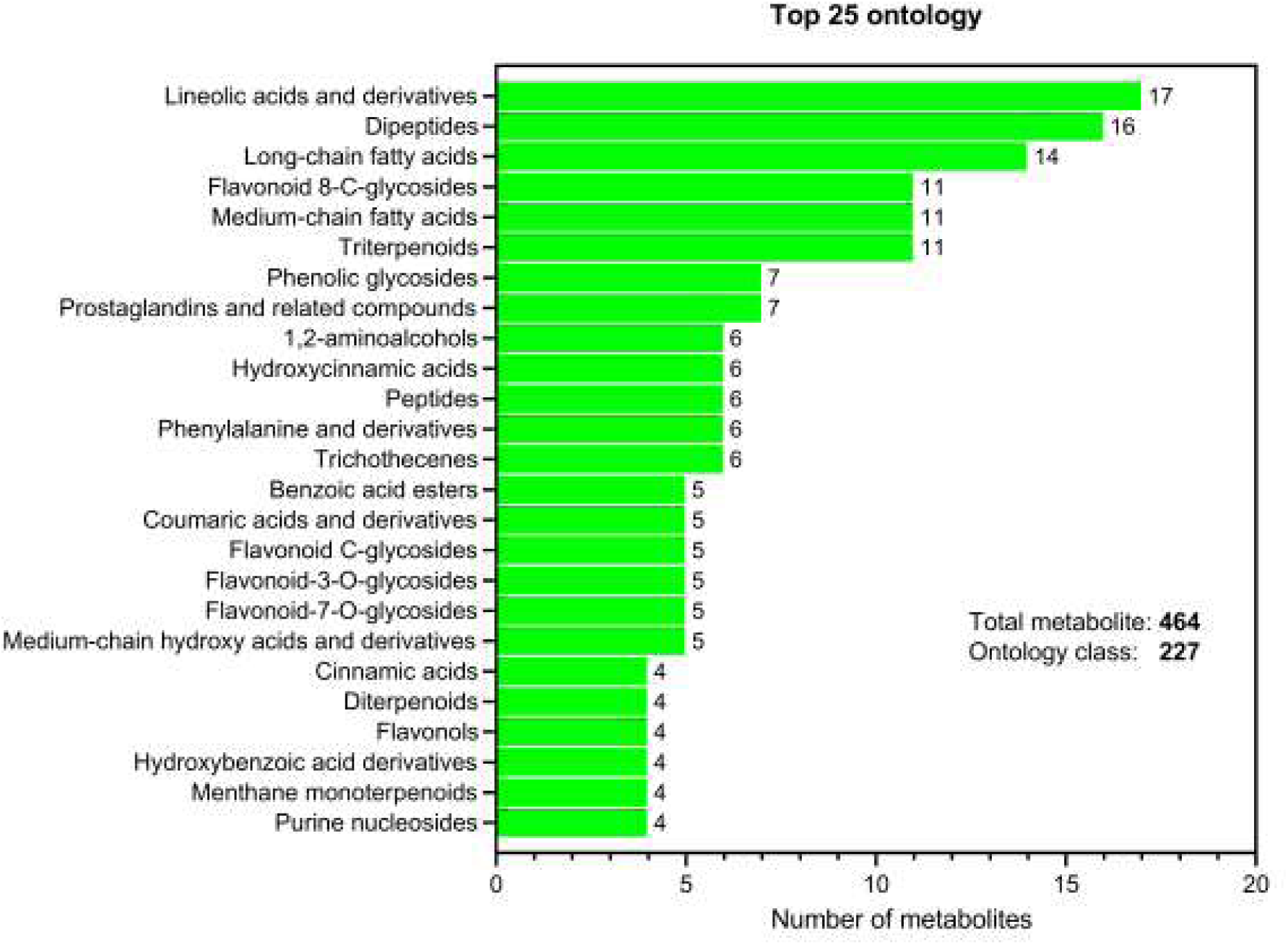
Untargeted LC–QTOF–MS/MS-based metabolomic profiling of *W. globosa*– derived nanovesicles. The horizontal bar chart illustrates the top 25 metabolite ontology classes identified through untargeted LC–QTOF–MS/MS. A total of 464 metabolites were annotated across 227 ontology classes. The number of annotated metabolites within each ontology class is indicated at the end of each bar. Full list of identified metabolites is provided in Supplementary File 2.

The HPLC-based validation further provided additional evidence supporting the LC-QTOF-MS/MS metabolomic findings by comparing the chromatographic profiles of WGNVs with those of selected reference compounds, including isovitexin, orientin, isoorientin, and schaftoside. The reference standards exhibited distinct retention times, allowing their chromatographic behavior to be compared with peaks detected in the WGNV sample (Figure 6 and Supplementary Figure 1). The WGNV chromatogram showed several peaks within the retention-time range of these standards, suggesting the presence of multiple chromatographically distinguishable constituents. These findings support the metabolomic results highlighting flavonoid glycosides as important constituents of *W. globosa*-derived nanovesicles.

**Figure 6.**
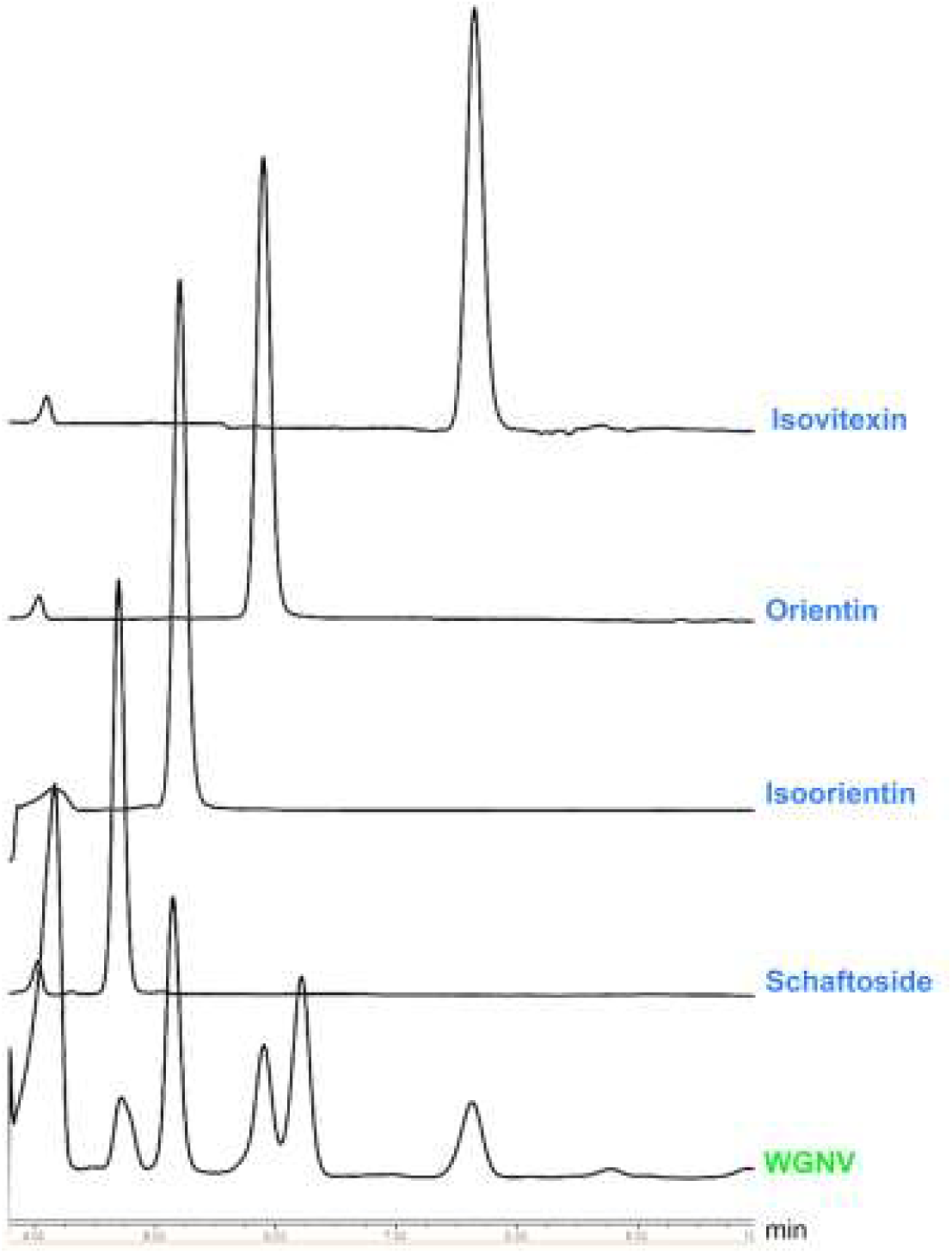
HPLC chromatograms of WGNV, schaftoside, isoorientin, orientin, and isovitexin detected at 254 nm. The standard compound peaks were detected at 4.7, 5.1, 5.9, and 7.6 min.

### 3.5 The internalization of WGNVs in neuronal cells demonstrated the inter-kingdom communication

Next, the bioactivities of WGNVs were investigated in an *in vitro* preclinical model of neurodegeneration induced by methylglyoxal (MGX), a reactive metabolite associated with diabetes, aging, and neurodegenerative disorders [15,19,23,24,38,39]. The mouse catecholaminergic CAD cell line was used as a neuronal model because it is derived from the central nervous system and is well established for neurobiological research [26,40].

Owing to their characteristic lipid bilayer structure, plant-derived nanovesicles exhibit favorable biocompatibility, facilitating efficient cellular uptake and internalization by recipient mammalian cells. To investigate potential interkingdom interactions between WGNVs and the neuronal cells, native WGNVs were labeled with the lipophilic fluorescent dye PKH26 and co-incubated with mouse neuronal CAD cells. As shown in Figure 7a–b, punctate fluorescent signals (nanodot-like structures) were observed throughout CAD cells exposed to PKH26-labeled WGNVs, suggesting interactions between WGNVs and neuronal cells and supporting interkingdom communication between plant-derived nanovesicles and mammalian cells.

**Figure 7.**
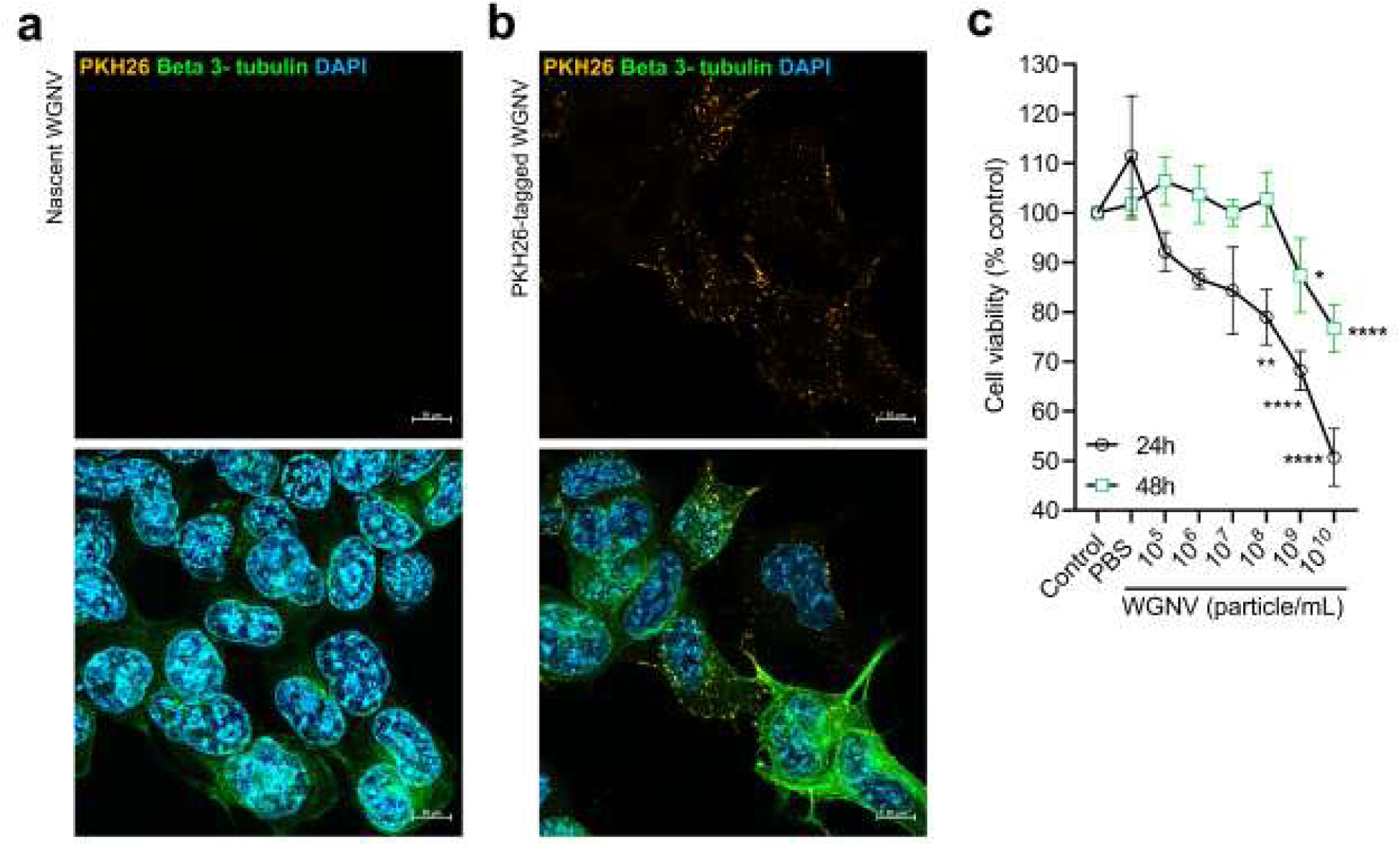
**Cellular internalization of *W.*** *globosa* derived nanovesicles and their concentration-dependent effects in the neuronal CAD cells. Representative fluorescence microscopy images of neuronal CAD cells exposed to **(a)** native (unlabeled) WGNVs or **(b)** PKH26-labeled WGNVs to evaluate cellular uptake. Immunocytochemical staining for βIII-tubulin (neuronal marker) and DAPI nuclear counterstaining were performed to visualize neuronal cell morphology and nuclei, respectively. Cellular uptake of WGNVs is indicated by the presence of intracellular punctate fluorescence signals. Scale bar: 10 μm. **(c)** Cell viability was assessed using the MTT assay following exposure to different concentrations of WGNVs (particles/mL) for 24 and 48 h. Cell viability was normalized to untreated controls and expressed as a percentage. Data are presented as mean ± standard deviation (SD) from four biological replicates (n = 4). Statistical significance was determined using one-way ANOVA. *p < 0.05, **p < 0.01, ****p < 0.0001 (vs the untreated control).

Furthermore, the effects of WGNVs on neuronal CAD cell viability were evaluated following exposure to WGNVs at concentrations ranging from 10^5^ – 10^10^ particle/mL for 24 and 48 h. A significant reduction in cell viability was observed after 24 h of exposure to WGNVs at 10^8^ particles/mL, with viability decreasing to approximately 80% of the control (Figure 7c). At 10^10^ particles/mL, cell viability further declined to approximately 50%. Therefore, a concentration below 10^7^ particles/mL was considered as the sublethal dose for subsequent investigations of the neuroprotective properties of *W. globosa*–derived nanovesicles.

### 3.6 WGNVs attenuated methylglyoxal-induced oxidative stress response, highlighting their biological relevance to neuroprotection

To evaluate the neuroprotective potential of WGNVs, MGX was used as a stress inducer associated with aging- and diabetes-related neurodegeneration. MTT assay was performed to establish the dose–response relationship between MGX concentration and neuronal CAD cell viability and to select an appropriate concentration for subsequent experiments. MGX significantly reduced CAD cell viability at concentrations ranging from 50 to 400 μM, with cell viability declining to below 20% at concentrations above 400 μM (Figure 8a).

**Figure 8.**
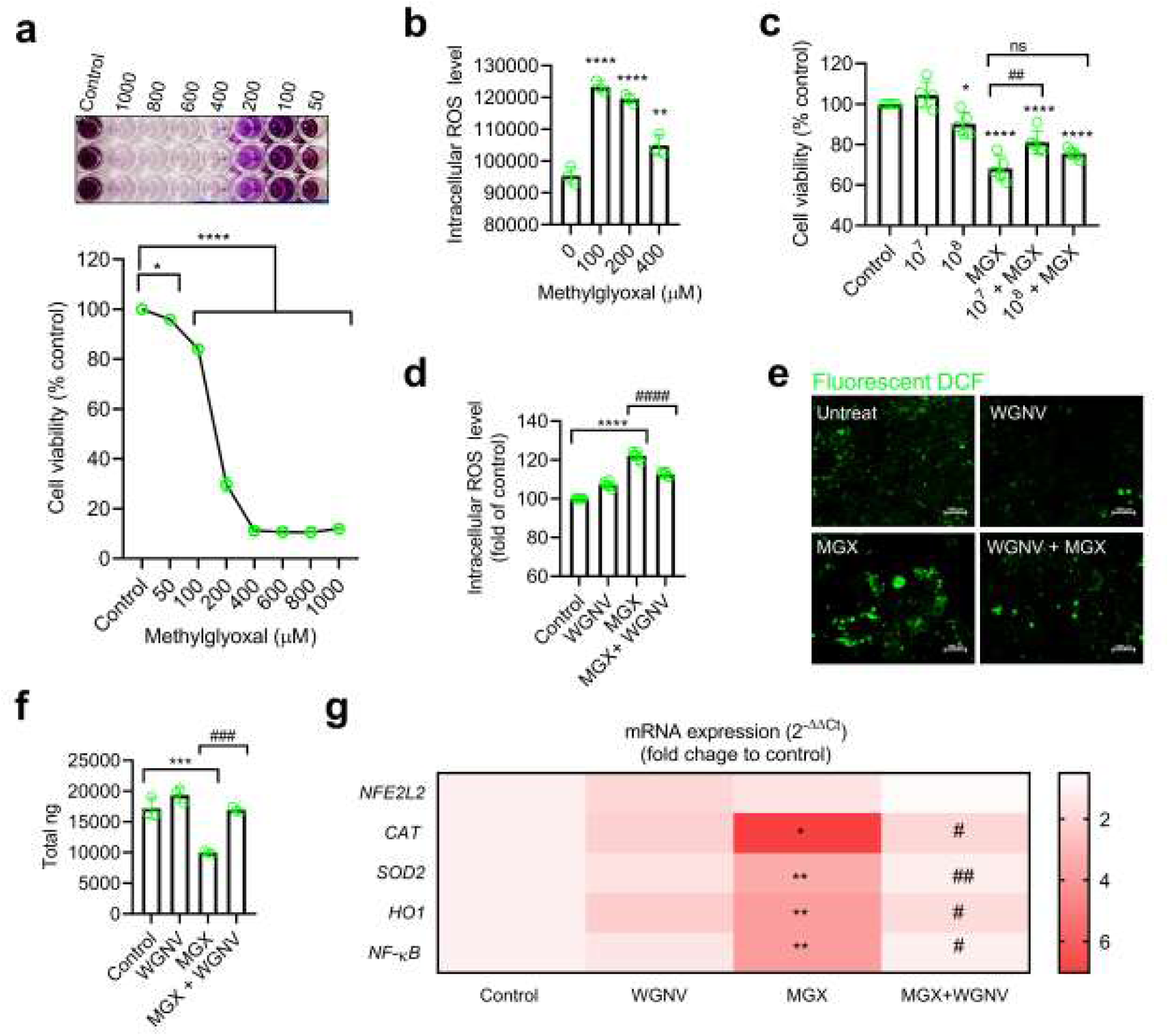
**Neuroprotective effects of *W.*** *globosa*–derived nanovesicles (WGNVs) against methylglyoxal (MGX)-induced oxidative stress response in CAD cells. **(a)** Representative images of formazan crystals and quantitative analysis of cell viability assessed by the MTT assay following treatment with different concentrations of MGX (50–1,000 μM) for 48 h. Absorbance was measured at 570 nm. **(b)** Quantification of intracellular reactive oxygen species (ROS) levels using the DCFH-DA assay following treatment with MGX (100–400 μM) for 4 h. **(c)** Quantitative analysis of cell viability assessed by the MTT assay in cells pretreated with WGNVs (10^7^ or 10^8^particles/mL) for 24 h, followed by exposure to MGX (100 μM) for an additional 24 h. **(d)** Quantification of intracellular ROS levels in cells pretreated with WGNVs (10^7^ particles/mL) for 24 h, followed by exposure to MGX (100 μM) for an additional 4 h. **(e)** Representative fluorescence images showing intracellular ROS levels, indicated by dichlorofluorescein (DCF) fluorescence, under the treatment conditions described in (d). **(f)** Total RNA yield (ng) extracted from cells subjected to the treatment conditions described in (c). **(g)** Heatmap showing the relative mRNA expression levels of oxidative stress-responsive genes in CAD cells subjected to the treatment conditions described in (f). Data are presented as mean ± SEM, with each dot representing an individual biological replicate. Experiments were performed using at least three biological replicates. Statistical significance was determined using one-way ANOVA. *p < 0.05, **p < 0.01, and ****p < 0.0001 (vs the control group). #p < 0.05, ##p < 0.01, ###p < 0.001, and ####p < 0.0001 (vs the MGX-treated group).

MGX is known to induce intracellular ROS accumulation, contributing to oxidative stress. To investigate this effect in neuronal CAD cells, cells were exposed to MGX at concentrations ranging from 100 to 400 μM, and intracellular ROS levels were measured after 4 h of treatment using the DCFH-DA fluorescent probe. MGX significantly increased ROS-associated fluorescence at all tested concentrations, indicating elevated intracellular oxidative stress (Figure 8b). The highest fluorescence intensity was observed at 100 μM MGX, which was therefore selected for subsequent experiments.

To evaluate the cytoprotective effects of WGNVs against MGX-induced stress in neuronal CAD cells, cells were pretreated with WGNVs at concentrations of 10^7^ or 10^8^ particles/mL for 24 h before exposure to MGX (100 μM) for an additional 24 h. Cell viability assays showed that pretreatment with WGNVs at the sublethal concentration of 10^7^ particles/mL protected neuronal CAD cells against MGX-induced reductions in cell viability (Figure 8c). In contrast, WGNVs at 10^8^ particles/mL did not confer cytoprotection, potentially due to their concentration-dependent cytotoxic effects, as previously demonstrated in Figure 7c. The restoration of cell viability was accompanied by a reduction in intracellular ROS levels, indicating that WGNV pretreatment attenuated MGX-induced ROS accumulation in neuronal CAD cells (Figure 7d–e). Together, the DPPH, FRAP, and cellular ROS assays support the antioxidant potential of WGNVs, demonstrated by their antioxidant activity in chemical assays and their ability to reduce intracellular ROS in neuronal CAD cells.

To further investigate the molecular responses associated with the cytoprotective effects of WGNVs, we evaluated changes in the expression of genes involved in oxidative stress responses following MGX exposure and WGNV treatment. As shown in Figure 8f, MGX exposure resulted in a reduction in total RNA yield compared with the control, whereas WGNV treatment was associated with a restoration of RNA yield toward control levels. Despite the absence of substantial cell death following MGX exposure, as indicated by the cell viability assay, MGX treatment induced pronounced upregulation of antioxidant-related genes (*Cat*, *Sod2*, and *Ho1*) and the inflammation-associated transcription factor gene *Nfkb2* (Figure 8g). These transcriptional alterations may represent an adaptive cellular response to MGX-induced oxidative stress, reflecting changes in antioxidant defense and inflammation-associated gene expression. Interestingly, WGNV treatment significantly attenuated the MGX-induced upregulation of these genes, suggesting its potential to modulate transcriptional responses associated with oxidative stress. Together with improved cell viability, these findings provide further evidence supporting the cytoprotective potential of WGNVs against MGX-induced neuronal stress.

### 3.7 WGNVs mitigated MGX-induced morphological alterations and preserved neuronal cellular integrity

To further investigate the cytoprotective potential of WGNVs, we examined morphological alterations in neuronal CAD cells following MGX exposure using βIII-tubulin immunofluorescence staining. As shown in Figure 9a, MGX-treated cells exhibited a more compact morphology, characterized by reduced cellular spreading and fewer apparent cellular extensions compared with control cells. Interestingly, WGNV treatment appeared to preserve cellular spreading and a more extended morphology following MGX exposure. To quantitatively assess these morphological alterations, cell area, roundness, and solidity were analyzed. MGX exposure significantly reduced cell area compared with the control (Figure 9b), whereas WGNV treatment showed a trend toward restoring cell area, although the difference compared with MGX treatment did not reach statistical significance (p = 0.0689). Additionally, MGX-treated cells exhibited a tendency toward increased roundness, consistent with the more compact and rounded appearance observed in the microscopy images. Although these differences were not statistically significant, WGNV treatment showed a trend toward reducing cell roundness, suggesting a partial shift toward the morphology observed in control cells (Figure 9c). Furthermore, MGX treatment significantly increased cellular solidity, indicating a shift toward a more compact and less morphologically complex phenotype. Notably, WGNV treatment significantly reduced MGX-induced increases in solidity, restoring values toward control levels (Figure 9d). Collectively, these findings suggest that WGNVs attenuated MGX-induced morphological alterations, particularly by preserving cellular spreading and reducing morphological compactness. These observations further support the potential of WGNVs to maintain neuronal morphology and cytoskeletal organization under MGX-induced stress conditions.

**Figure 9.**
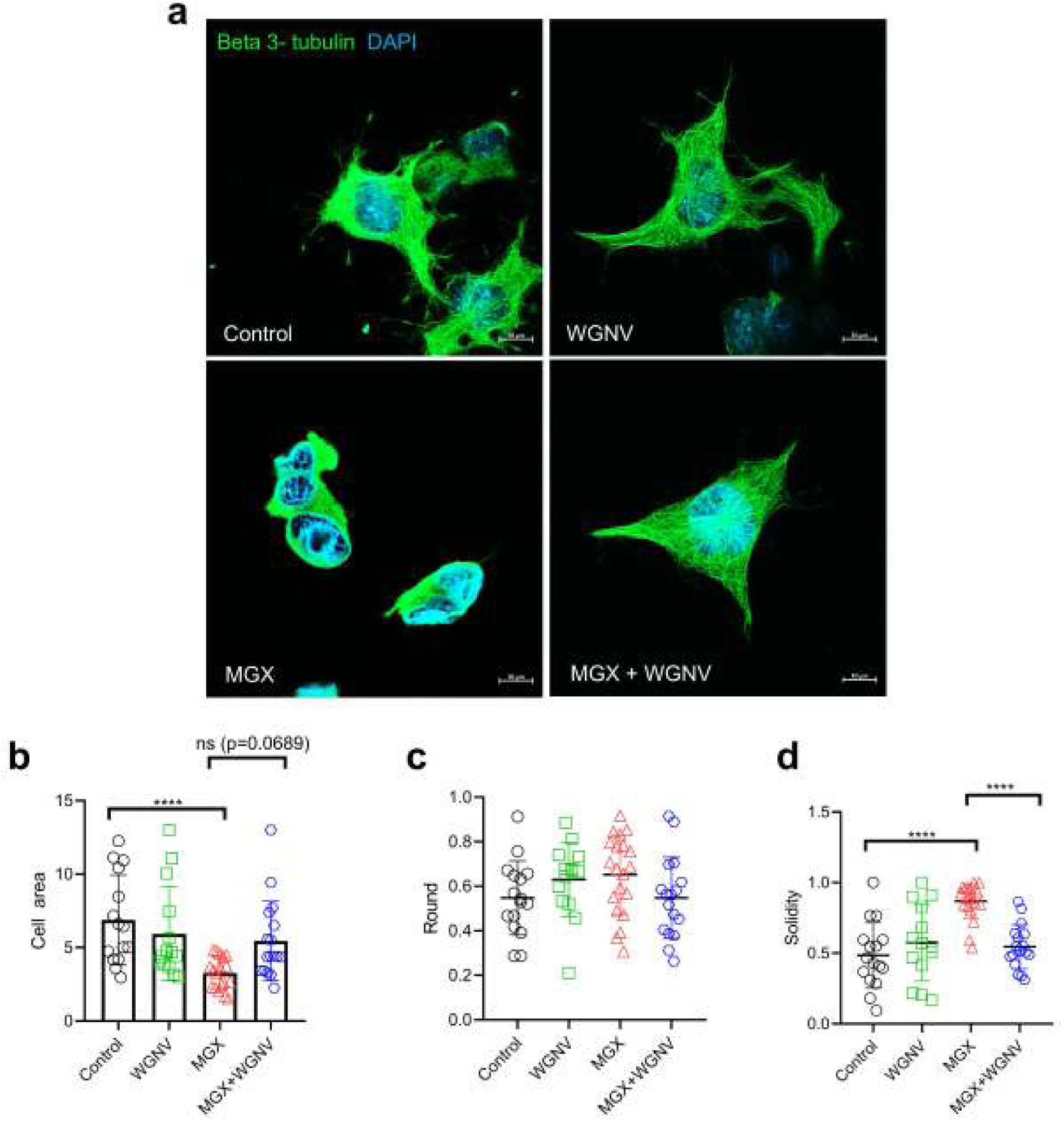
WGNVs preserved cell shape integrity against MGX-induced morphological alterations in CAD neuronal cells. **(a)** Representative confocal images showing cell morphology and the microtubule cytoskeleton in CAD neuronal cells following different treatments. Cells were pretreated with WGNVs (10^7^ particles/mL) for 24 h, followed by exposure to MGX (100 μM) for 4 h. Cells were then fixed, permeabilized, and stained for beta-3-tubulin (green), a neuron-specific microtubule marker, and DAPI (blue) to visualize nuclei. Scale bar is 10μm. Quantification of cell shape parameters, including **(b)** cell area (μm^2^), **(c)** roundness, and **(d)** solidity. Each dot represents an individual cell analyzed across three biological replicates. Statistical significance among experimental groups was determined using one-way ANOVA followed by multiple-comparisons test. ****p < 0.0001 in the respective compared groups.

## 4. Discussion

This study highlights the potential of *Wolffia globosa*, an emerging future food, as a source of bioactive exosome-like nanovesicles that exhibit antioxidant activity and protect neuronal cells against methylglyoxal-induced oxidative stress in vitro. By integrating nanovesicle characterization, metabolite profiling, and functional evaluation, this work provides a foundation for exploring the transition from conventional functional food resources to plant-derived nanomedicine platforms.

Plant-derived nanovesicles (PDNV) have emerged as a promising research area in nanomedicine because of their potential to carry bioactive compounds and interact with mammalian cells. Research in this field has exponentially increased[5,6,11,41,42]. In the present study, we investigated *W. globosa*–derived nanovesicles (WGNVs) isolated and purified using a combination of ultracentrifugation and ultrafiltration. Nanoparticle tracking analysis (NTA), zeta potential measurements, and cryogenic electron microscopy (cryo-EM) were used to characterize their physical properties and structural features. NTA revealed a modal particle size of 127.5 nm and a mean particle size of 187 nm, while cryo-EM visualized a double-membrane structure. The recovered WGNV sample reached a particle concentration of approximately 10^11^ particles/mL, demonstrating the ability of the isolation workflow to yield a concentrated nanovesicle preparation. Previous studies have reported two approaches for isolating WGNVs. Gradient-based enrichment yielded particles with a mean size of 105.3 nm and a concentration on the order of 10^10^ particles/mL [13], whereas differential ultracentrifugation produced nanovesicles ranging from 94.6-106.9 nm[14]. Differences in isolation procedures, sample processing, and particle characterization may account for the variation in reported particle sizes and concentrations. However, these findings demonstrate that *W. globosa* is a promising source of PDNVs for investigating their bioactive composition and potential applications in nanomedicine.

Despite these advances in isolation and physicochemical characterization, biochemical markers that could support the identification of WGNVs and other PDNVs remain insufficiently established. This presents a significant challenge compared with mammalian extracellular vesicles, for which commonly used protein markers, such as CD9, CD63, and CD81, are available[25,43]. Given the considerable biochemical diversity among plant species, identifying candidate biochemical markers specific to PDNVs is an important research priority. Addressing this gap could improve the identification, characterization, and comparability of plant-derived nanovesicle preparations across studies.

Although a growing body of evidence has reported antioxidant activity in *W. globosa* biomass and crude extracts, the antioxidant properties of WGNVs remain unexplored. [2,3,12,44]. A comprehensive literature search identified two recent publications reporting the isolation of WGNVs. One focused primarily on developing an isolation protocol[13], whereas the other explored their biological application as a doxorubicin delivery platform for cancer treatment[14]. However, neither study systematically characterized the metabolite composition or overall phytochemical profile of the isolated WGNVs. We therefore addressed this knowledge gap. The WGNV samples exhibited antioxidant activity in both DPPH radical-scavenging and ferric reducing antioxidant power (FRAP) assays, alongside detectable total phenolic and flavonoid contents, supporting their potential antioxidant capacity. To our knowledge, the present study provides the first evidence of antioxidant properties in WGNVs.

The present study also employed an untargeted, unbiased metabolomics approach to comprehensively profile the metabolites associated with WGNVs. Major metabolite classes detected were amino acids and dipeptides, purine/pyrimidine nucleosides, phenolic and hydroxycinnamic acids, organic acids, coumarins, flavonoids, fatty acids, and sphingolipids (Supplementary File 2). Flavonoids represented the most abundant and diverse class of metabolites detected in WGNVs, including apigenin and its derivatives, such as vitexin, isovitexin, isoschaftoside, vicenin II, and apigenin-6-C-glucoside-7-O-glucoside. Other identified flavonoids included luteolin glycosides, such as orientin and isoorientin, as well as kaempferol and quercetin derivatives. HPLC validation confirmed the presence of selected metabolites, including isovitexin, orientin, isoorientin, and schaftoside, supporting the reliability of the untargeted LC–QTOF– MS/MS metabolomics approach. Additionally, the detection of sphingolipids, including sphingosine, sphinganine, and C17 sphinganine, provides further insight into the lipid composition of the WGNV preparation. These findings support the presence of diverse plant-associated metabolites in the isolated nanovesicles, although additional analyses are required to establish the specificity of these compounds as markers of plant-derived nanovesicles.

Given the antioxidant activity and diverse bioactive metabolites identified in WGNVs, we further investigated the biological activities of the isolated nanovesicles. Neurodegenerative diseases represent a growing public health challenge with their prevalence increasing alongside population aging. Their multifactorial pathogenesis and the limited availability of disease-modifying treatments highlight the need to explore novel preventive and therapeutic strategies. In this context, PDNVs have attracted increasing interest as potential bioactive platforms. WGNVs therefore represent a promising candidate for further investigation, particularly in the context of developing sustainable, food-derived nanovesicle platforms for neuroprotection. Methylglyoxal (MGX) is a reactive byproduct of glycolysis that can accumulate under conditions associated with diabetes, aging, and neurodegenerative diseases[15,16], where it may contribute to neuronal dysfunction through oxidative stress and cellular damage[19,21,39,45]. Several phytochemicals have been investigated for their protective effects against MGX-induced neuronal damage[16,22,24,46]. However, to our knowledge, no previous studies have reported the neuroprotective effects of PDNVs against MGX-induced neuronal degeneration. In the present study, MGX exposure reduced neuronal cell viability, increased oxidative stress and altered the expression of stress-related genes, and disrupted cellular integrity in the mouse CNS-derived CAD cell line. WGNV treatment mitigated these adverse effects, suggesting its potential to protect neuronal cells against MGX-induced oxidative stress response.

Notably, WGNVs were internalized by the neuronal cells, supporting their potential for cellular delivery and further investigation as a platform for transporting plant-derived bioactive compounds. However, cellular internalization alone does not establish their ability to cross the blood–brain barrier (BBB), which remains a major challenge for delivering therapeutic agents to the CNS. Future in vio studies should therefore investigate the biodistribution of WGNVs following different administration routes and determine whether they can reach brain tissue. Intranasal administration is an emerging route for delivering mammalian extracellular vesicles to the central nervous system, but the feasibility of this approach for plant-derived nanovesicles requires further investigation[47–50]. In addition, the antioxidant and neuroprotective effects of WGNVs should be evaluated in vivo using relevant models of neurological disorders, including Alzheimer’s disease (AD).

AD is a progressive neurodegenerative disorder and the most common cause of dementia[51]. Although current treatments do not cure the disease, acetylcholinesterase (AChE) inhibition is an established therapeutic strategy for managing its cognitive symptoms[52,53]. AChE catalyzes the breakdown of acetylcholine (ACh), a neurotransmitter essential for learning and memory. Previous studies have reported the AChE-inhibitory potential of stem cell-derived exosomes[33]. However, the potential of PDNVs to modulate AChE activity remains largely unexplored. In addition to their neuroprotective effects against MGX-induced oxidative stress, our preliminary findings demonstrated that *W. globosa*–derived nanovesicles exhibited AChE-inhibitory activity in vitro (Supplementary Figure 3). These findings highlight WGNVs as a promising candidate for further investigation of their potential relevance to AD-related research.

## 5. Conclusion

The recent study demonstrated the successful isolation and characterization of nanovesicles from *Wolffia globosa*, an emerging future functional food with promising nutritional and biomedical potential. WGNVs exhibited a nanoscale size distribution and a lipid bilayer structure, possessed intrinsic antioxidant activity, and contained a diverse repertoire of bioactive metabolites, particularly phenolic and flavonoid compounds. Of particular relevance, WGNV treatment attenuated MGX-induced ROS accumulation, modulated stress-responsive gene expression, and mitigated neuronal morphological alterations, supporting the neuroprotective potential of WGNVs. A visual summary of this study is presented in Figure 10. Collectively, these findings highlight the opportunity to harness nanovesicles from the future functional foods as naturally derived bioactive platforms. Further investigations may broaden the potential applications of WGNVs in neuroprotection and neurodegenerative disease research, including Alzheimer’s disease, as well as other biomedical fields, such as gut–brain axis modulation, colitis, and biocosmetic applications.

**Figure 10.**
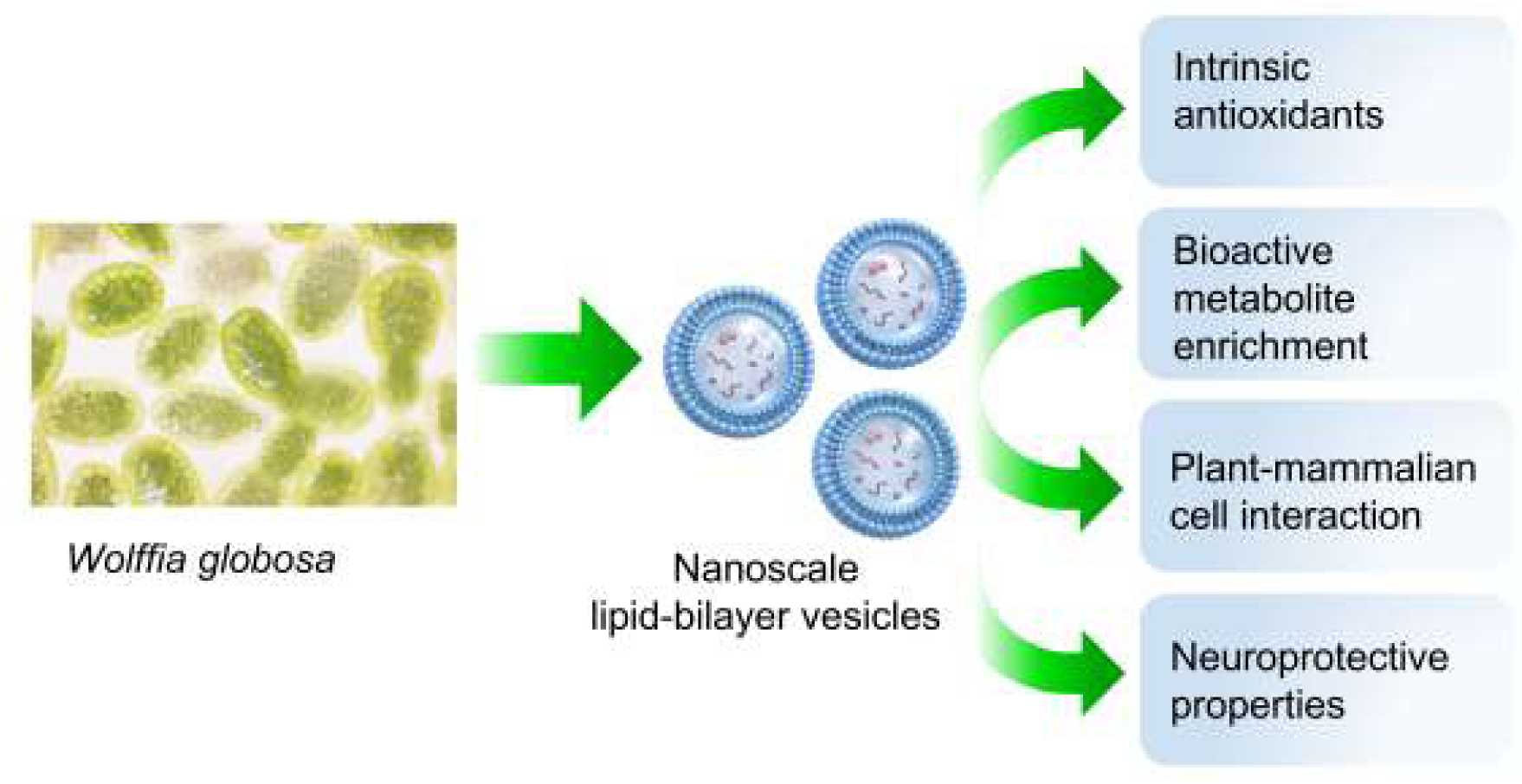
A visual summary of this study. The isolation of *W. globosa*–derived nanovesicles and the characterization of their antioxidant capacity, bioactive metabolite composition, cellular internalization, and neuroprotective potential against methylglyoxal-induced oxidative stress highlight their promise for advancing conventional functional foods toward nanomedicine and expanding their utility across a broad range of biomedical applications.

## Declaration of Generative AI and AI-assisted technologies in the writing process

The author used ChatGPT to assist with English language editing and grammatical improvements during the preparation of this manuscript. Following the use of this tool, the author carefully reviewed and revised the content as necessary and takes full responsibility for the content of the published article.

## Contributions (CRediT)

EK: Conceptualization, Data curation, Formal analysis, Investigation, Methodology, Validation, Visualization, Writing – original draft, Funding acquisition.

PB: Data curation, Formal analysis, Investigation, Methodology, Validation, Visualization, Writing – original draft.

MS and TC (contributed equally): Data curation, Formal analysis, Investigation, Methodology

## Acknowledgment

The authors sincerely thank the Frontier Research Facility (FRF), Mahidol University, for providing cryogenic electron microscopy (cryo-EM) and zeta potential analysis services. The authors also acknowledge the Central Instrument Facility, Institute of Molecular Biosciences, Mahidol University, for providing access to high-resolution confocal microscopy and nanoparticle tracking analysis (NTA). Special thanks are extended to Pipob Suwanchaikasem of Baiya Phytopharm Co., Ltd., Bangkok, Thailand, for providing metabolomics services. We also thank Tharita Kitisripanya and Waraporn Putalun of the Faculty of Pharmaceutical Sciences, Khon Kaen University, Khon Kaen, Thailand, for providing standard compounds for metabolite validation, and Benyakan Pongkitwitoon of the Department of Pharmaceutical Botany, Faculty of Pharmacy, Mahidol University, Bangkok, Thailand, for providing access to HPLC-grade solvents.

## Data availability

All data supporting the findings of this study are available from the corresponding author, Dr. Ekkaphot Khongkla, upon reasonable request.

## References

[1] G. Jahreis, K.S. Sree, C. Dawczynski, K.J. Appenroth, Water lentil rich in nutrients and bioactive components-A powerful group of vegetables for humanity’s future plant-based diet, Trends in Food Science & Technology 170 (2026) 105600. 10.1016/j.tifs.2026.105600.

[2] N.K. Yadav, A.B. Patel, S. Debbarma, M.B. Priyadarshini, G. Kumar, S. Baidya, A.D. Upadhyay, Characterization of phenolic compounds in watermeal (Wolffia globosa) through LC-ESI-QTOF-MS/MS: assessment of bioactive compounds, in vitro antioxidant and anti-diabetic activities following different drying methods, Food Measure 18 (2024) 8651–8672. 10.1007/s11694-024-02833-y.

[3] R. Khonkarn, K. Daowtak, O. Kraseasintra, T. Luetragoon, K. Usuwanthim, K. Taynawa, K. Chanphong, Bioactive Potential of Protein Extracts Derived from Dried Wolffia globosa on In Vitro Antioxidant Activities and Pro-Inflammatory Cytokine Production, Molecules 30 (2025) 4092. 10.3390/molecules30204092.

[4] A.E. Hernani-Rodrigo, S.F. Waters, L.S.V. Figueroa, J.G. Msemakweli, N. LaRuta-Rosas, M. Buhezo-Chamon, R.H. Gilman, Duckweed (Wolffia globosa, Wolffia arrhiza, and Lemna minor) as novel therapeutic food for acute malnutrition and famine alleviation in low- and middle-income countries and humanitarian crisis settings, Front. Nutr. 13 (2026). 10.3389/fnut.2026.1835959.

[5] S. Du, D. Jia, G. Liang, Y. Dou, Plant-derived nanovesicles: the intelligent nanoplatforms for therapeutics and drug delivery, J Nanobiotechnol 24 (2026) 582. 10.1186/s12951-026-04484-1.

[6] C.K. Belkare, O.V. Daware, Plant-derived exosome-like nanovesicles as programmable bio-organic excipients: a next-generation platform for precision and immuno-silent drug delivery, DARU J Pharm Sci 34 (2026) 35. 10.1007/s40199-026-00617-x.

[7] K.-J. Lo, M.-H. Wang, C.-T. Ho, M.-H. Pan, Plant-Derived Extracellular Vesicles: A New Revolutionization of Modern Healthy Diets and Biomedical Applications, J. Agric. Food Chem. 72 (2024) 2853–2878. 10.1021/acs.jafc.3c06867.

[8] Y. Zhu, J. Zhao, H. Ding, M. Qiu, L. Xue, D. Ge, G. Wen, H. Ren, P. Li, J. Wang, Applications of plant-derived extracellular vesicles in medicine, MedComm 5 (2024) e741. 10.1002/mco2.741.

[9] M.D. Langellotto, G. Rassu, C. Serri, S. Demartis, P. Giunchedi, E. Gavini, Plant-derived extracellular vesicles: a synergetic combination of a drug delivery system and a source of natural bioactive compounds, Drug Deliv. and Transl. Res. 15 (2025) 831–845. 10.1007/s13346-024-01698-4.

[10] M. Liu, H. Wang, Y. Yang, J. Wang, Z. Sun, S. Zhao, W. Kong, W. Sun, C. Wang, L. Feng, Therapeutic potential of plant-derived exosome-like nanoparticles for CNS diseases: advancements in preparation and characterization, Annals of Medicine 57 (2025) 2537345. 10.1080/07853890.2025.2537345.

[11] Md. M.N. Azim, Y.K. Lee, M.J. Alam, J.-H. Kim, J.H. Park, Therapeutic potential of plant-derived exosome-like nanovesicles as a phytomedicine in age-related diseases, Biomedicine & Pharmacotherapy 199 (2026) 119441. 10.1016/j.biopha.2026.119441.

[12] N.K. Yadav, A.B. Patel, S. Debbarma, M.B. Priyadarshini, H. Priyadarshi, Characterization of Bioactive Metabolites and Antioxidant Activities in Solid and Liquid Fractions of Fresh Duckweed (*Wolffia globosa*) Subjected to Different Cell Wall Rupture Methods, ACS Omega 9 (2024) 19940–19955. 10.1021/acsomega.3c09674.

[13] A. Jiraviriyakul, M. Piamano, K. Pimpa, N. Sangprasert, M. Wongchaiya, S. Arin, C. Nensat, W. Songjang, Enrichment and Characterization Protocol for Plant-Derived Nanovesicles from the Aquatic Plant Wolffia globosa, Current Protocols 6 (2026) e70427. 10.1002/cpz1.70427.

[14] C. Nensat, W. Songjang, M. Piamano, K. Pimpa, N. Sangprasert, C. Srisuwan, A. Jiraviriyakul, RGD-functionalized Wolffia globosa-derived Exosome-like Nanoparticles for Targeted Doxorubicin Delivery in Breast Cancer Therapy, Pharm Nanotechnol (2026). 10.2174/01221173854464260828044240.

[15] C.G. Schalkwijk, C.D.A. Stehouwer, Methylglyoxal, a Highly Reactive Dicarbonyl Compound, in Diabetes, Its Vascular Complications, and Other Age-Related Diseases, Physiological Reviews 100 (2020) 407–461. 10.1152/physrev.00001.2019.

[16] R. Kold-Christensen, M. Johannsen, Methylglyoxal Metabolism and Aging-Related Disease: Moving from Correlation toward Causation, Trends in Endocrinology & Metabolism 31 (2020) 81–92. 10.1016/j.tem.2019.10.003.

[17] A. Croce, P. Mirra, F.C. Pignalosa, F. Zatterale, I. Prevenzano, D. Conza, L. Ulianich, C. Miele, F. Beguinot, F. Fiory, Methylglyoxal Affects Dopamine Homeostasis in SH-SY5Y Cells Through the Modulation of miR-190a and miR-214, The FASEB Journal 40 (2026) e71639. 10.1096/fj.202501805R.

[18] H. Wang, S. Boeren, W. Bakker, I.M.C.M. Rietjens, E. Saccenti, L. Zheng, An integrated proteomics and metabolomics analysis of methylglyoxal-induced neurotoxicity in a human neuroblastoma cell line, Npj Sci Food 8 (2024) 84. 10.1038/s41538-024-00328-0.

[19] H.J. Chun, Y. Lee, A.H. Kim, J. Lee, Methylglyoxal Causes Cell Death in Neural Progenitor Cells and Impairs Adult Hippocampal Neurogenesis, Neurotox Res 29 (2016) 419–431. 10.1007/s12640-015-9588-y.

[20] J.M.T. Chu, D.K.M. Lee, D.P.K. Wong, G.T.C. Wong, K.K.M. Yue, Methylglyoxal-induced neuroinflammatory response in in vitro astrocytic cultures and hippocampus of experimental animals, Metab Brain Dis 31 (2016) 1055–1064. 10.1007/s11011-016-9849-3.

[21] S. Di Loreto, V. Caracciolo, S. Colafarina, P. Sebastiani, A. Gasbarri, F. Amicarelli, Methylglyoxal induces oxidative stress-dependent cell injury and up-regulation of interleukin-1β and nerve growth factor in cultured hippocampal neuronal cells, Brain Research 1006 (2004) 157–167. 10.1016/j.brainres.2004.01.066.

[22] M.R. de Oliveira, I.C. Custódio de Souza, C.R. Fürstenau, Promotion of mitochondrial protection by naringenin in methylglyoxal-treated SH-SY5Y cells: Involvement of the Nrf2/GSH axis, Chem Biol Interact 310 (2019) 108728. 10.1016/j.cbi.2019.108728.

[23] C. Angeloni, M. Malaguti, B. Rizzo, M.C. Barbalace, D. Fabbri, S. Hrelia, Neuroprotective Effect of Sulforaphane against Methylglyoxal Cytotoxicity, Chem. Res. Toxicol. 28 (2015) 1234–1245. 10.1021/acs.chemrestox.5b00067.

[24] Y.-J. Chen, X.-B. Huang, Z.-X. Li, L.-L. Yin, W.-Q. Chen, L. Li, Tenuigenin protects cultured hippocampal neurons against methylglyoxal-induced neurotoxicity, European Journal of Pharmacology 645 (2010) 1–8. 10.1016/j.ejphar.2010.06.034.

[25] E. Khongkla, K. Promtap, J. Meerasri, P. Mo-Mai, W. Chankamngoen, N. Sirinonthanawech, B. Chetsawang, Human adipose stem cell-derived exosomes modulate the transcriptome of D-galactose-induced neuronal cells, Sci Rep 16 (2026) 4575. 10.1038/s41598-025-34661-5.

[26] E. Khongkla, K. Uppakara, N. Boonmuen, K. Bhukhai, W. Saengsawang, A Novel Methodology Using Dexamethasone to Induce Neuronal Differentiation in the CNS-Derived Catecholaminergic CAD Cells, Cell Mol Neurobiol 42 (2022) 2337–2353. 10.1007/s10571-021-01109-z.

[27] E. Khongkla, P. Boonsnongcheep, P. Suwanchaikasem, K. Promthep, M. Srisaisup, T. Charuraksa, P. Makarathut, B. Chetsawang, First evidence for isolation and characterization of Bacopa monnieri (L.) Wettst-derived nanovesicles with anti-neuroblastoma potential, Sci Rep (2026). 10.1038/s41598-026-55427-7.

[28] G. Midekessa, K. Godakumara, J. Ord, J. Viil, F. Lättekivi, K. Dissanayake, S. Kopanchuk, A. Rinken, A. Andronowska, S. Bhattacharjee, T. Rinken, A. Fazeli, Zeta Potential of Extracellular Vesicles: Toward Understanding the Attributes that Determine Colloidal Stability, ACS Omega 5 (2020) 16701–16710. 10.1021/acsomega.0c01582.

[29] I.F.F. Benzie, J.J. Strain, The Ferric Reducing Ability of Plasma (FRAP) as a Measure of “Antioxidant Power”: The FRAP Assay, Analytical Biochemistry 239 (1996) 70–76. 10.1006/abio.1996.0292.

[30] A. Nicolescu, C.I. Bunea, A. Mocan, Total flavonoid content revised: An overview of past, present, and future determinations in phytochemical analysis, Analytical Biochemistry 700 (2025) 115794. 10.1016/j.ab.2025.115794.

[31] P. Boonsnongcheep, P. Pichetpongtorn, P. Juntosuth, T. Yowsong, P. Vansanit, S. Preeprame, T. Kanchanapoom, W. Putalun, LED and methyl jasmonate enhance a strong antioxidant compound, verbascoside, production in Clerodendrum indicum and Acanthus ebracteatus, J. Plant Biochem. Biotechnol. 31 (2022) 98–106. 10.1007/s13562-021-00659-z.

[32] E.A. Ainsworth, K.M. Gillespie, Estimation of total phenolic content and other oxidation substrates in plant tissues using Folin–Ciocalteu reagent, Nat Protoc 2 (2007) 875–877. 10.1038/nprot.2007.102.

[33] M.I. Khan, E.S. Jeong, M.Z. Khan, J.H. Shin, J.D. Kim, Stem cells-derived exosomes alleviate neurodegeneration and Alzheimer’s pathogenesis by ameliorating neuroinflamation, and regulating the associated molecular pathways, Sci Rep 13 (2023) 15731. 10.1038/s41598-023-42485-4.

[34] H. Tsugawa, T. Cajka, T. Kind, Y. Ma, B. Higgins, K. Ikeda, M. Kanazawa, J. VanderGheynst, O. Fiehn, M. Arita, MS-DIAL: data-independent MS/MS deconvolution for comprehensive metabolome analysis, Nat Methods 12 (2015) 523–526. 10.1038/nmeth.3393.

[35] S. Lee, S. Hwang, M. Seo, K.B. Shin, K.H. Kim, G.W. Park, J.Y. Kim, J.S. Yoo, K.T. No, BMDMS-NP: A comprehensive ESI-MS/MS spectral library of natural compounds, Phytochemistry 177 (2020) 112427. 10.1016/j.phytochem.2020.112427.

[36] P. Ghensi, E. Bressan, C. Gardin, L. Ferroni, M.C. Soldini, F. Mandelli, C. Soldini, B. Zavan, The Biological Properties of OGI Surfaces Positively Act on Osteogenic and Angiogenic Commitment of Mesenchymal Stem Cells, Materials 10 (2017) 1321. 10.3390/ma10111321.

[37] H. Kim, X. Xue, Detection of Total Reactive Oxygen Species in Adherent Cells by 2’,7’-Dichlorodihydrofluorescein Diacetate Staining, Journal of Visualized Experiments (JoVE) (2020) e60682. 10.3791/60682.

[38] J. Lu, E. Randell, Y. Han, K. Adeli, J. Krahn, Q.H. Meng, Increased plasma methylglyoxal level, inflammation, and vascular endothelial dysfunction in diabetic nephropathy, Clinical Biochemistry 44 (2011) 307–311. 10.1016/j.clinbiochem.2010.11.004.

[39] S. Dukic-Stefanovic, R. Schinzel, P. Riederer, G. Münch, AGES in brain ageing: AGE-inhibitors as neuroprotective and anti-dementia drugs?, Biogerontology 2 (2001) 19–34. 10.1023/a:1010052800347.

[40] E. Khongkla, J. Kwanthongdee, B. Chetsawang, Proteomic profiling reveals mitochondrial metabolic alterations in dexamethasone-induced neuronal differentiation, Science Progress 109 (2026) 00368504261459907. 10.1177/00368504261459907.

[41] Y. Huang, S. Wang, Q. Cai, H. Jin, Effective methods for isolation and purification of extracellular vesicles from plants, JIPB 63 (2021) 2020–2030. 10.1111/jipb.13181.

[42] R. Wang, Y. Zhang, Y. Guo, W. Zeng, J. Li, J. Wu, N. Li, A. Zhu, J. Li, L. Di, P. Cao, Plant-derived nanovesicles: Promising therapeutics and drug delivery nanoplatforms for brain disorders, Fundamental Research 5 (2025) 830–850. 10.1016/j.fmre.2023.09.007.

[43] J.A. Welsh, D.C.I. Goberdhan, L. O’Driscoll, E.I. Buzas, C. Blenkiron, B. Bussolati, H. Cai, D. Di Vizio, T.A.P. Driedonks, U. Erdbrügger, J.M. Falcon-Perez, Q.-L. Fu, A.F. Hill, M. Lenassi, S.K. Lim, M.G. Mahoney, S. Mohanty, A. Möller, R. Nieuwland, T. Ochiya, S. Sahoo, A.C. Torrecilhas, L. Zheng, A. Zijlstra, S. Abuelreich, R. Bagabas, P. Bergese, E.M. Bridges, M. Brucale, D. Burger, R.P. Carney, E. Cocucci, R. Crescitelli, E. Hanser, A.L. Harris, N.J. Haughey, A. Hendrix, A.R. Ivanov, T. Jovanovic-Talisman, N.A. Kruh-Garcia, V. Ku’ulei-Lyn Faustino, D. Kyburz, C. Lässer, K.M. Lennon, J. Lötvall, A.L. Maddox, E.S. Martens-Uzunova, R.R. Mizenko, L.A. Newman, A. Ridolfi, E. Rohde, T. Rojalin, A. Rowland, A. Saftics, U.S. Sandau, J.A. Saugstad, F. Shekari, S. Swift, D. Ter-Ovanesyan, J.P. Tosar, Z. Useckaite, F. Valle, Z. Varga, E. van der Pol, M.J.C. van Herwijnen, M.H.M. Wauben, A.M. Wehman, S. Williams, A. Zendrini, A.J. Zimmerman, M. Consortium, C. Théry, K.W. Witwer, Minimal information for studies of extracellular vesicles (MISEV2023): From basic to advanced approaches, Journal of Extracellular Vesicles 13 (2024) e12404. 10.1002/jev2.12404.

[44] P. Temviriyanukul, W. Inthachat, T. Laowanitwattana, P. Buacheen, U. Suttisansanee, P. Pitchakarn, Optimization and Metabolomic Profiling of Wolffia globosa Extract Reveal Multitarget Neuroprotective Activity Against Alzheimer’s Disease with In Vitro, In Silico, and In Vivo Validation, International Journal of Molecular Sciences 27 (2026) 8266. 10.3390/ijms27188266.

[45] S. Victor-Sami, A. Kamali-Roosta, Y.A. Shamsaldeen, Methylglyoxal induces death in human brain neuronal cells (SH-SY5Y), prevented by metformin and dapagliflozin, Journal of Diabetes and Its Complications 38 (2024) 108832. 10.1016/j.jdiacomp.2024.108832.

[46] J.M.T. Chu, D.K.M. Lee, D.P.K. Wong, R.N.S. Wong, K.K.L. Yung, C.H.K. Cheng, K.K.M. Yue, Ginsenosides attenuate methylglyoxal-induced impairment of insulin signaling and subsequent apoptosis in primary astrocytes, Neuropharmacology 85 (2014) 215–223. 10.1016/j.neuropharm.2014.05.029.

[47] M. Umezawa, F. Suyama, K. Tachibana, A. Onoda, K. Takeda, Plant-Derived Nanovesicle Enhanced Microribonucleic Acid (MicroRNA) Transfer from Nasal Cavity to the Brain, Mol. Pharmaceutics 22 (2025) 6964–6971. 10.1021/acs.molpharmaceut.5c00931.

[48] P. Li, S. Sun, X. Zhu, X. Liu, R. Yin, Y. Chen, J. Chang, L. Ye, J. Gao, X. Zhao, H. Xu, Y. Wang, W. Zuo, Z. Sun, S. Wang, X. Zhang, J. Wei, R.C. Zhao, Q. Han, Intranasal delivery of engineered extracellular vesicles promotes neurofunctional recovery in traumatic brain injury, J Nanobiotechnol 23 (2025) 229. 10.1186/s12951-025-03181-9.

[49] L.D. Moss, D. Sode, R. Patel, A. Lui, C. Hudson, N.A. Patel, P.C. Bickford, Intranasal delivery of exosomes from human adipose derived stem cells at forty-eight hours post injury reduces motor and cognitive impairments following traumatic brain injury, Neurochemistry International 150 (2021) 105173. 10.1016/j.neuint.2021.105173.

[50] L.N. Madhu, M. Kodali, S. Rao, S. Attaluri, R. Upadhya, G. Shankar, B. Shuai, Y. Somayaji, S.V. Ganesh, V.S. Kumar, J.E. James, P.A. Shetty, A. LeMaire, X. Rao, J.J. Cai, A.K. Shetty, Intranasal Human NSC-Derived EVs Therapy Can Restrain Inflammatory Microglial Transcriptome, and NLRP3 and cGAS-STING Signalling, in Aged Hippocampus, Journal of Extracellular Vesicles 15 (2026) e70232. 10.1002/jev2.70232.

[51] J. Zhang, Y. Zhang, J. Wang, Y. Xia, J. Zhang, L. Chen, Recent advances in Alzheimer’s disease: mechanisms, clinical trials and new drug development strategies, Sig Transduct Target Ther 9 (2024) 211. 10.1038/s41392-024-01911-3.

[52] N.V. Mehta, A. Kapadia, M. Khambete, A. Abhyankar, The Evolution of AChE Inhibitors in Alzheimer’s Disease: From Single-Target to Multi-Target Ligands, ChemBioChem 26 (2025) e202500053. 10.1002/cbic.202500053.

[53] H. Cheng, L. Kang, Y. Xu, R.E. Tanzi, C. Zhang, C. Wang, Innovation breakthrough in the Alzheimer’s disease pharmaceutical industry, Npj Drug Discov. 3 (2026) 9. 10.1038/s44386-026-00044-7.

