## Supplementary Information for "*Wolffia globosa*–derived nanovesicles as a natural source of bioactive metabolites with neuroprotective potential against methylglyoxal-induced oxidative stress"

**Supplemented Figure 1.** Chromatographic profiles obtained in **(a)** positive and **(b)** negative ionization modes, comparing the blank and WGNV samples.

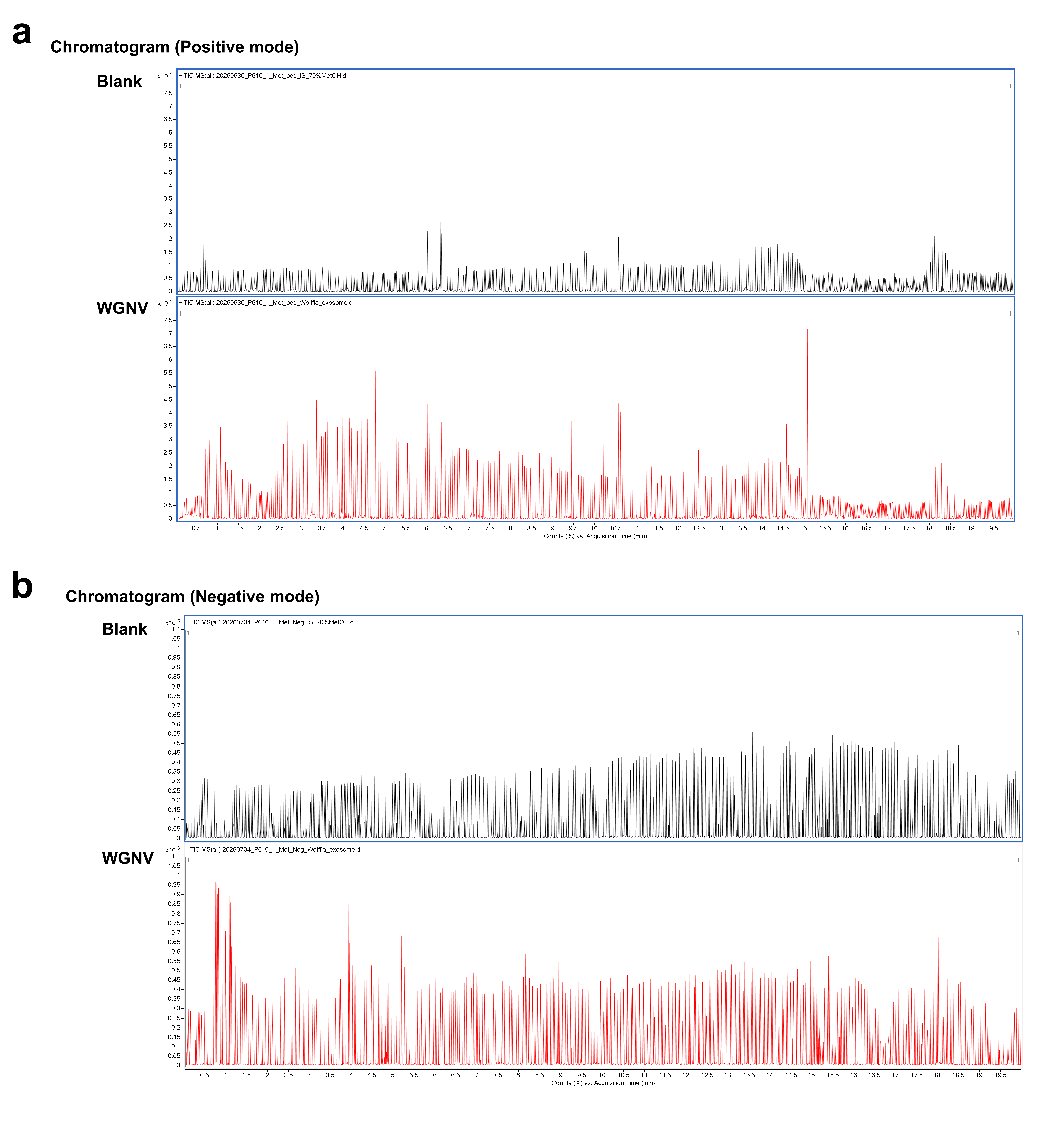

**Supplemented Figure 2.** UV spectra of peaks detected in WGNV sample (A1, B1, C1, and D1) compare with the UV spectra of standard compounds. (A2) schaftoside, (B2) isoorientin, (C2) orientin, and (D2) isoorientin.

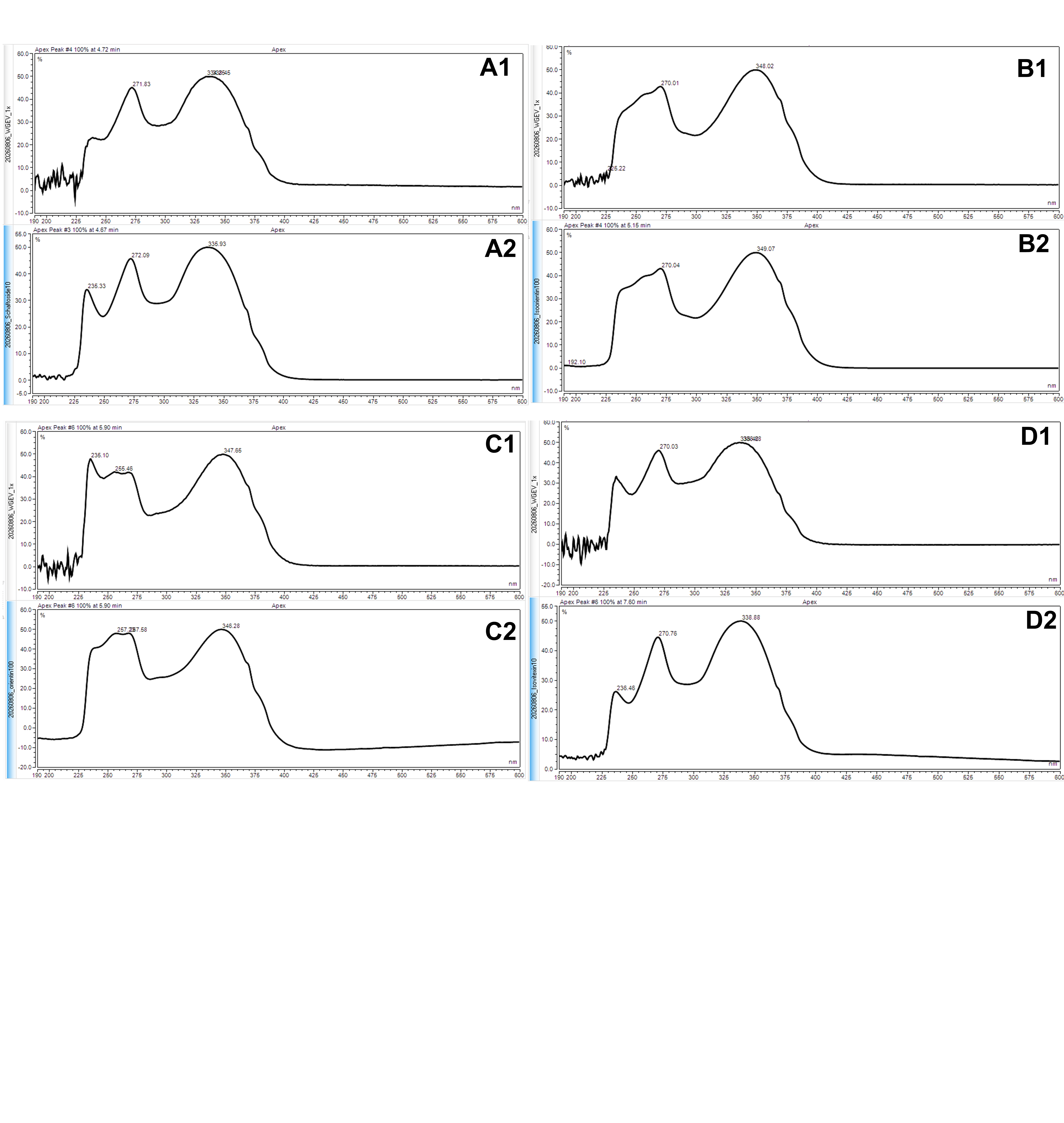

**Supplemented Figure 3. (a)** Schematic diagram illustrating the experimental protocol used to evaluate the acetylcholinesterase (AChE) inhibitory potential of *Wolffia globosa*–derived nanovesicles (WGNVs) in human neuroblastoma SH-SY5Y cells. **(b)** Relative AChE activity (%) in SH-SY5Y cells following treatment with WGNVs (10^7^ particles/mL), expressed relative to the control. Statistical significance was determined using Student’s t-test. **** p < 0.0001 (n = 4).

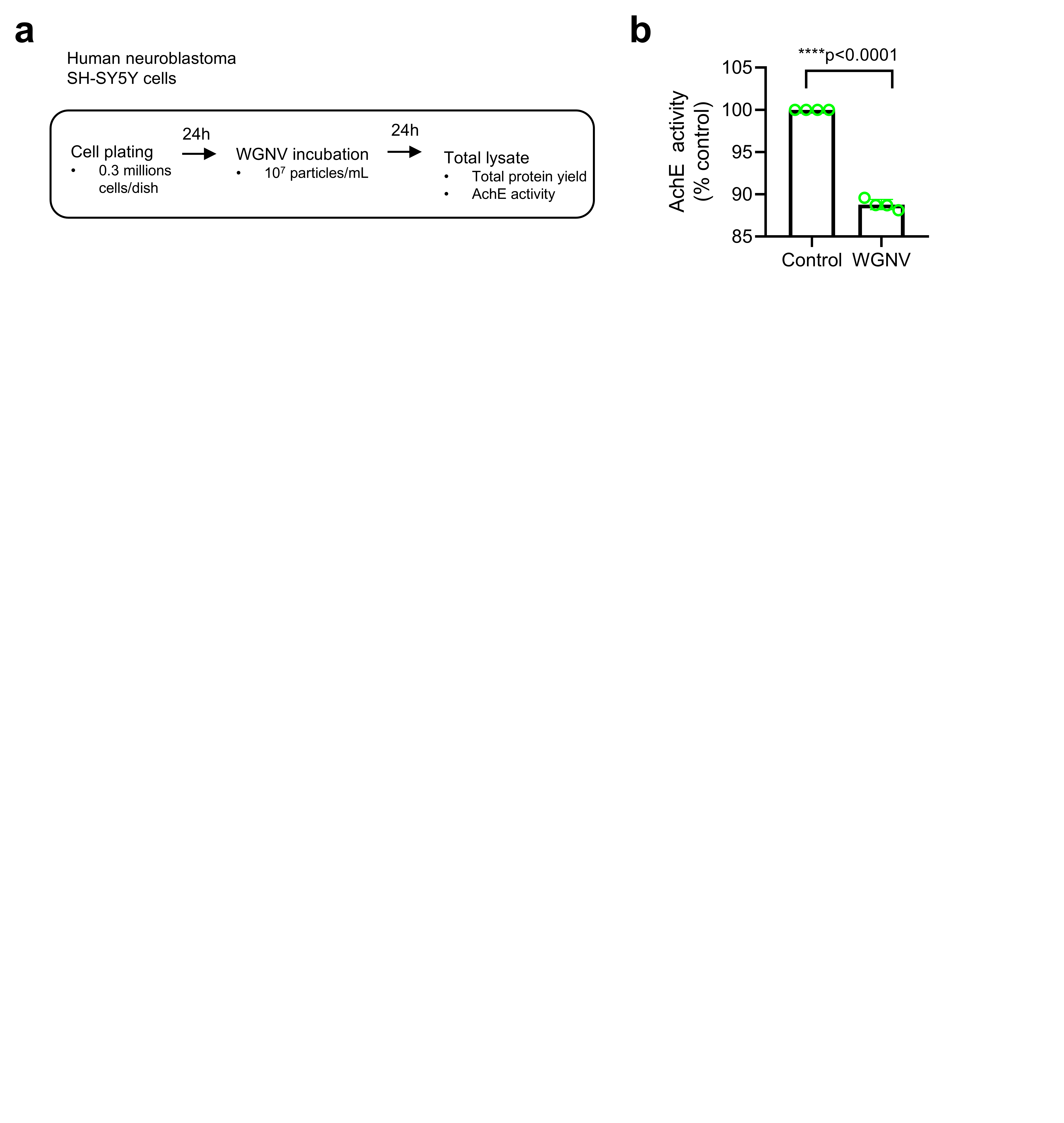

**Supplemented File 1.** Representative NTA video showing the Brownian motion of WGNVs during nanoparticle tracking analysis.

**Supplemented File 2.**  List of putatively identified metabolites from untargeted LC–QTOF–MS/MS-based metabolomic analysis of WGNVs in negative and positive ionization modes.
